# The type 1 diabetes gene *TYK2* regulates β-cell development and its responses to interferon-α

**DOI:** 10.1101/2022.02.22.481272

**Authors:** Vikash Chandra, Hazem Ibrahim, Clémentine Halliez, Rashmi Prasad, Federica Vecchio, Om Prakash Dwivedi, Jouni Kvist, Diego Balboa, Jonna Saarimäki-Vire, Hossam Montaser, Tom Barsby, Väinö Lithovius, Isabella Artner, Swetha Gopalakrishnan, Leif Groop, Roberto Mallone, Decio L. Eizirik, Timo Otonkoski

## Abstract

Type 1 diabetes (T1D) is an autoimmune disease that results in the destruction of insulin producing pancreatic β-cells. One of the genes associated with T1D is *TYK2,* which encodes a Janus kinase with critical roles in type-Ι interferon (IFN) mediated intracellular signaling. To study the role of TYK2 in human pancreatic β-cell development and response to IFNα, we generated *TYK2* knockout human iPSCs and directed them into the pancreatic endocrine lineage. Here we show that loss of TYK2 compromised the emergence of endocrine precursors by regulating KRAS expression while mature stem cell-islets (SC-islets) function was not affected. In the maturing SC-islets, the loss or inhibition of TYK2 prevented IFNα-induced antigen processing and presentation, including MHC Class Ι expression in pancreatic endocrine and progenitor cells. Furthermore, in a CD8^+^ cytotoxic T-cell co-culture model, the survival of β-cells was enhanced by a selective TYK2 inhibitor. These results identify an unsuspected role for TYK2 on β-cell development and support TYK2 inhibition in adult β-cells as a potent therapeutic target to halt T1D progression.

## Introduction

Type 1 diabetes (T1D) is a chronic autoimmune disease characterized by pancreatic islet inflammation resulting in eventual specific loss of the insulin-secreting β-cells(1, 2). Genome-wide association and other genetic studies have identified more than 120 non-HLA regions associated with the risk of developing T1D(3). One such predisposing gene is tyrosine kinase 2 (*TYK2*), a member of the JAK (Janus Kinase) family, which plays a critical role in intracellular signal transducer and activator of transcription (STAT) signaling stimulated by cytokines, including type I interferons (IFN-Ι)(4). IFN-I signaling is involved in the aetiology of T1D through up-regulation of MHC Class Ι expression and antigen presentation, which leads to the targeting of cytotoxic autoimmunity towards β-cells(5, 6). A recent report suggests that rare loss-of-function *TYK2* promoter mutations are associated with increased diabetes susceptibility in the Japanese population(7, 8). Intriguingly, some of the non-synonymous single-nucleotide polymorphisms (SNPs) of *TYK2* (rs34536443 and rs2304256) that induce a partial inhibition of *TYK2* expression are associated with protection against several autoimmune diseases, including T1D and rheumatoid arthritis(9, 10). As the majority of the T1D candidate genes are expressed in human islets and regulate their function(2), it is important to decipher their role in pancreatic development, function and responses to immune challenges, so that stage specific therapeutic interventions can be effectively developed. Human pluripotent stem cells (hPSCs) and their efficient differentiation towards pancreatic stem cell-islets (SC-islets) provide preeminent tools for studying human pancreatic development and the role for candidate genes acting at the β-cell level(11–14).

We presently studied the expression of T1D candidate genes in the developing endocrine pancreas and identified *TYK2* as one of the upregulated genes. The effects of TYK2 perturbation were studied by single-cell transcriptomics in the SC-islets throughout their development. Surprisingly, specific *TYK2* knockout (KO) stem cell lines showed compromised endocrine precursor formation, whereas TYK2 inhibition in differentiated islet cells completely blocked the IFNα responses. Based on our results, TYK2 inhibition of mature islet cells is a potential interventional therapy for T1D.

## Results

### Expression of T1D candidate genes during pancreatic differentiation

To identify candidate genes of pancreatic development that are also associated with the development of T1D in humans, we performed deep RNA sequencing (∼ 200 million reads per sample) at specific stages of human induced pluripotent stem cells (hiPSCs) pancreatic differentiation (Figure 1A). Our differentiation strategy robustly phenocopied human pancreatic development as revealed by the expression pattern of key genes of pancreatic developmental hierarchy (Figure 1B). Screening of stage-specific transcriptome revealed differential expression of several T1D candidate genes, e.g., *TYK2*, *DLK1* and *IFIH1* were strongly upregulated while *PRKCQ*, *BACH2* and *GPR183* were downregulated as differentiation progressed towards endocrine lineage (Figure 1C). The significant increase of TYK2 expression (*p*=2.4x10^-12^) was further confirmed at transcript and protein level in independent experiments (Figure 1, D and E). In agreement with this observation, RNAseq data on human fetal pancreas of mid-to-late 1^st^ trimester (n=16) showed a strong up-regulation of *TYK2* transcription with increasing gestational age (Figure 1F).

**Figure 1.**
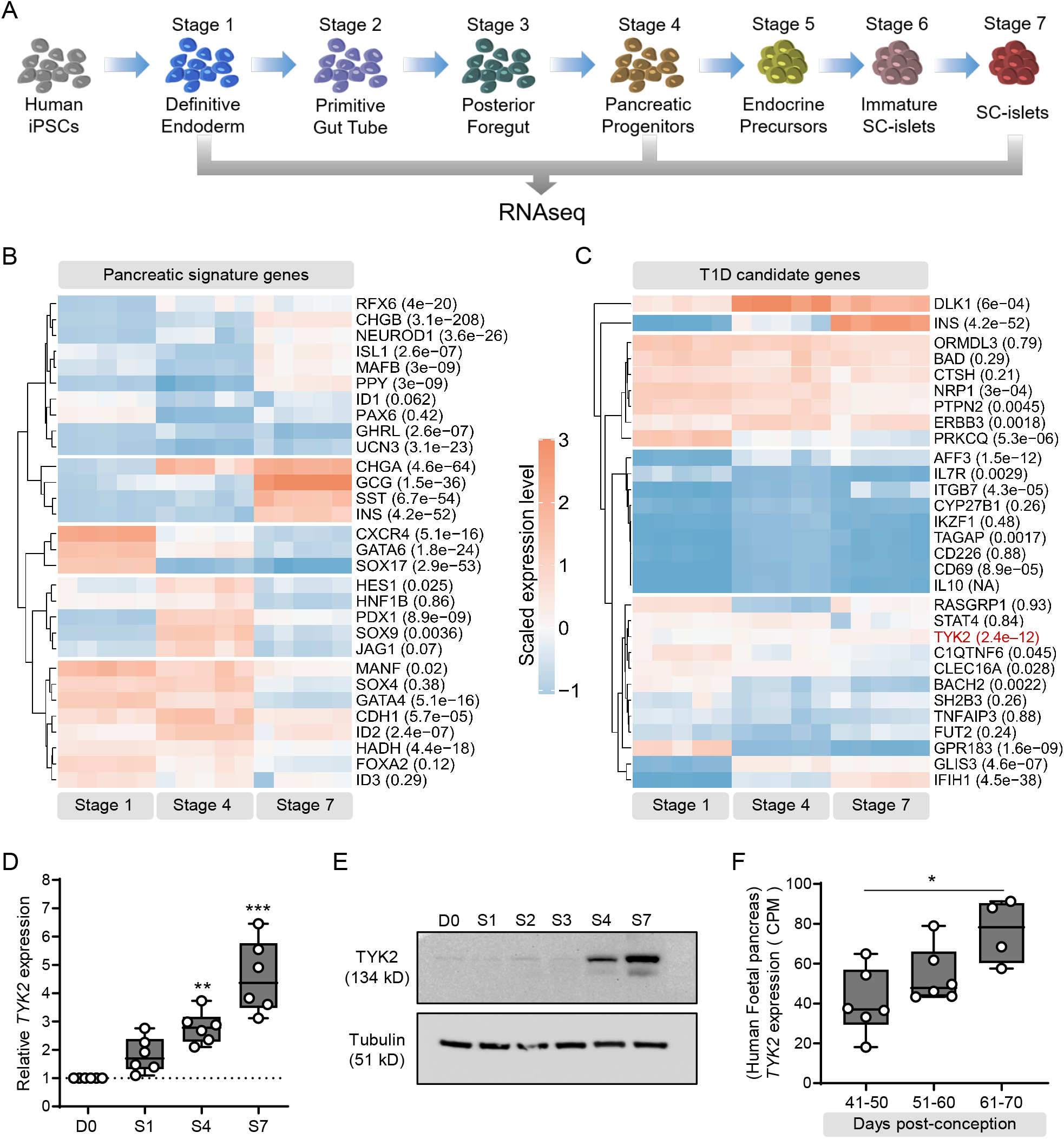
Expression pattern of *TYK2*, a T1D candidate gene, during human pancreatic development. **(A)** Schematic representation of the pancreatic differentiation protocol. **(B)** Heatmap of pancreatic lineage genes from deep RNAseq analysis at Stage 1 (definitive endoderm), Stage 4 (pancreatic-progenitors) and Stage 7 (SC-islets). **(C)** Similar heatmap for known T1D candidate genes. Each gene is shown with a multiple testing corrected *p* value generated for the longitudinal differential expression of the gene during differentiation (n = 5). **(D)** Expression of *TYK2* during pancreatic differentiation shown by qRT-PCR for the transcripts (n = 6) and **(E)** immunoblot analysis for the expression of TYK2 protein. Tubulin was used as a loading control (n = 3). **(F)** Expression pattern of *TYK2* in human fetal pancreas samples at 40 to 70 days post conception (n = 16). For **D** and **F**, significance was determined using one-way ANOVA with Tukey’s multiple comparison test. *p < 0.05; **p < 0.01; ***p < 0.001.

### Loss or inhibition of TYK2 results in defective endocrine precursor (EP) formation

Next, we employed CRISPR-Cas9 to generate *TYK2* KO hiPSC lines (HEL46.11)(12). Two KO clones, selected by the loss of TYK2 protein expression, were used in this study and a non-edited negative clone served as a wildtype (WT) control (Supplemental Figure 1, A-C). Sanger sequencing confirmed the precise deletion of the targeted 277 bp sequence of the ATG-start codon-containing exon 3 (Supplemental Figure 1D) with no evidence of CRISPR-induced off-target indels. Bulk RNAseq based e-Karyotyping(15) and immunocytochemical analysis of WT and *TYK2* KO hiPSC lines confirmed the absence of genomic aberrations and showed similar expression levels for key proliferation and pluripotency markers (Supplemental Figure 1, E-H).

TYK2 is known to associate with IFN-Ι receptor (IFNAR1) but not IFN-ΙΙ receptor and activate downstream STAT signaling(16). Stimulation with IFNα activated STAT1 and STAT2 in the WT but not in the KO (Figure 2A). In contrast, IFNγ treatment activated STAT1 in both WT and *TYK2* KO cells, further confirming the specific inhibition of IFNAR1 signaling in *TYK2* KO cells.

**Figure 2.**
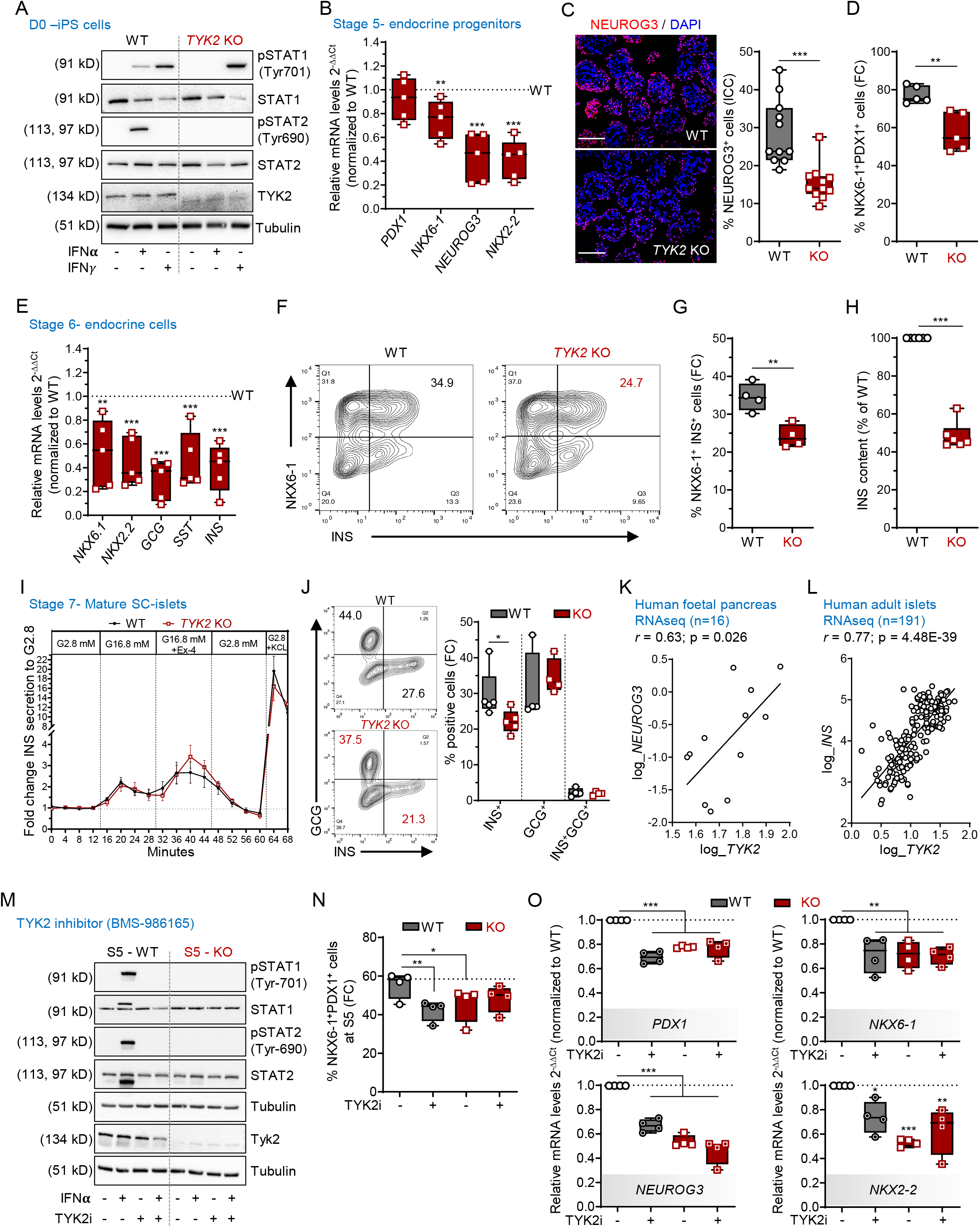
Loss of TYK2 is associated with defective formation of endocrine precursors. **(A)** Immunoblot analysis after 30 min exposure to IFNα or IFNγ showing the phosphorylation status of STAT1 and STAT2 in WT and *TYK2* KO hiPSCs (n = 3). **(B)** Relative transcript levels of *PDX1*, *NKX6-1*, *NEUROG3* and *NKX2-2* in *TYK2* KO Stage 5 endocrine progenitors analysed by qRT-PCR (n = 5). **(C)** Immunocytochemistry for NEUROG3 with a representative image and the quantification of 3 separate experiments (3-4 images per experiment were analysed). Scale bar = 150 μm. **(D)** Flow cytometry analysis of PDX1^+^ NKX6-1^+^ endocrine progenitors (n = 5). **(E)** Stage 6 immature endocrine cells shown for relative transcript levels of *NKX6-1*, *NKX2-2*, *GCG*, *SST* and *INS* analysed by qRT-PCR (n = 5). **(F)** Representative contour plot of the flow cytometry analysis for the expression of NKX6-1^+^ and INS^+^ cells (Stage 6). **(G)** Quantification of the data shown in F (n = 4). **(H)** Total cellular INS content normalized to DNA content in Stage 6 (n = 5). **(I)** Dynamic insulin secretion of perifused WT and *TYK2* KO SC-islets at Stage 7. The cells were stimulated with 16.8 mM glucose, 50 ng/ml exendin-4 (Ex4) and 30 mM KCl. Values are normalized to the average secretion during 2.8 mM glucose, the first 16 minutes of the test. One-way ANOVA with Welch’s correction (n = 3), **(J)** Representative contour plot of the flow cytometry analysis for the INS^+^ GCG^+^ and polyhormonal cells and their percentage quantification (n = 4). **(K)** Correlation of *TYK2* and *NEUROG3* expression in RNA-seq of human fetal pancreas samples. Pearson’s correlation after log normalization of counts (n = 16). **(L)** Similar analysis for *TYK2* and *INS* expression in adult human islet samples by RNA-seq. Pearson’s *r* correlation test (n = 191). **(M)** WT and *TYK2* KO cells were treated with the TYK2 inhibitor BMS-986165 during Stage 3 – 5 and analysed for the phosphorylation of STAT1 and STAT2 in the presence of IFNα (30 min) (n = 3). **(N)** Flow cytometric analysis of PDX1^+^ NKX6-1^+^ cells after treatment with TYK2i at Stage 5. One-way ANOVA with Dunnett’s multiple comparison test (n = 4). **(O)** Expression of *PDX1*, *NKX6-1*, *NEUROG3* and *NKX2-2* with qRT-PCR from the same experiments as in panel N. Ordinary one-way ANOVA with Tukey’s multiple comparison (n = 4). Two-tailed unpaired t-test was performed to determine the significance levels unless otherwise mentioned. Box and whiskers plots showing min to max with all the points. *p < 0.05; **p < 0.01; ***p < 0.001.

To study the role of TYK2 in pancreatic development, we performed pancreatic differentiation using a seven-stage protocol(11, 12). The WT and *TYK2* KO lines showed comparable differentiation capacity until stage (S) 4 pancreatic progenitors (PP), as demonstrated by similar proportions of cells expressing CXCR4 at S1 (definitive endoderm) and co-expressing PDX1 and NKX6-1 at S4 (Supplemental Figure 2, A and B). In addition, the expression levels of *SOX9*, *FOXA2*, *PTF1A* and *PDX1* were similar at S4. However, NKX6-1 expression was significantly reduced in the KO compared to their WT counterpart (Supplemental Figure 2C). Endocrine precursors (EP) induced at S5 of the differentiation protocol. EP formation was significantly compromised in *TYK2* KO, as shown by decreased expression of transcripts for *NKX6-1* (25±7% reduction), *NEUROG3* (56±9%) and *NKX2-2* (58±7%) (Figure 2B). The number of NEUROG3^+^ cells was also significantly reduced (46.3% reduction, *p*=0.0002) as evaluated by immunocytochemistry (Figure 2C). The proportion of PDX1^+^NKX6-1^+^ cells was significantly reduced as evaluated by flow cytometry (19±4% reduction, *p*=0.004) (Figure 2D). S6 is marked by a substantial increase of immature endocrine cells. As a consequence of impaired EP formation, we also observed a global decrease in the expression of the islet hormone transcripts *INS* (59±8% reduction), *GCG* (70±7%) and *SST* (54±10%) at S6 (Figure 2E). This observation was further confirmed by flow cytometry for INS^+^NKX6-1^+^ cells (29.8±2% reduction) and normalized total insulin content (51.4±2% reduction) (Figure 2, F-H).

Dynamic insulin secretion was studied after 3 weeks of S7 maturation. Interestingly, the responses of *TYK2* KO cells to high glucose, to the glucagon-like peptide-1 (GLP1) analogue exendin-4 and to K^+^-induced membrane depolarization were comparable to WT (Figure 2I). This data indicates that the functional capacity of the *TYK2* KO SC-islets remained intact despite their reduced number of mono-hormonal INS^+^ cells (Figure 2J). To investigate the *in vivo* functional potential of the SC-islets, we implanted equal numbers of WT and *TYK2* KO S7 SC-islets under the kidney capsule of non-diabetic NOD-SCID-Gamma mice (Supplemental Figure 2D). Blood glucose and circulating human C-peptide levels were measured until 2 months post-implantation. There was no significant difference between mice implanted with WT or *TYK2* KO SC-islets (Supplemental Figure 2, E and F). Correspondingly, we observed comparable proportions of mono-hormonal INS^+^ cells to total INS^+^ and GCG^+^ cells in WT and *TYK2* KO 2-month grafted tissue sections (Supplemental Figure 2, G and H). In line with this, the RNA-seq data from 16 human fetal pancreata also revealed a significant positive correlation between *TYK2* and *NEUROG3* (r=0.63; *p*=0.026; Figure 2K). Likewise, the adult islet RNA-seq dataset(17) (n=191) showed a strong positive correlation between *TYK2* and *INS* expression (r=0.77; *p*=4.4x10^-39^) (Figure 2L), suggesting a regulatory role of TYK2 in the pancreatic endocrine lineage.

These results indicate that the complete loss of TYK2 impairs EP formation but does not affect the subsequent maturation and function of β-cells. We next validated these findings by using a selective and potent allosteric TYK2 inhibitor (TYK2i) BMS-986165(18) during S3 to S5 differentiation. Upon IFNα stimulation of S5 WT cells, STAT1 and STAT2 activation were abolished by TYK2i (Figure 2M). We then investigated whether TYK2i could recapitulate the *TYK2* KO reduced EP formation phenotype. We observed a reduced number of PDX1^+^NKX6-1^+^ cells in TYK2i treated S5 WT cells (Figure 2N). Following TYK2i treatment at the end of S5, we also observed a similarly reduced expression of the transcripts *PDX1* (by 30±3%), *NEUROG3* (by 33±4%), *NKX6-1* (by 28±6%) and *NKX2-2* (by 26±8%) with no change in the TYK2i treated KO samples (Figure 2O). Notably, we also replicated the above findings of TYK2 inhibition on the H1 human embryonic stem cells (hESCs) (Supplemental Figure 3, A-E). Collectively, these data confirm that TYK2 regulates the formation of pancreatic EP.

### TYK2 negatively regulates KRAS expression

To understand the mechanism by which loss or inhibition of TYK2 compromises EP formation during pancreatic lineage differentiation, a whole transcriptome analysis was performed at the stem cell level (hiPSCs), S4 (PP), S5 (EP) and S6 (immature SC-islets) (Figure 3A). Principal component analysis (PCA) of respective transcriptomes for WT and KO genotype clustered together in order of developmental stage, but a significant difference (*p*=0.02) was observed at S5 (Figure 3B), with 319 up-regulated and 412 down-regulated genes (Figure 3C). In line with the findings described above, the key pancreatic transcription factors (TFs) *NEUROG3* and *NKX2-2* were significantly down-regulated at S5 (Figure 3, D and E). Reactome enrichment analysis confirmed the downregulation of gene sets associated with “Regulation of β-cell Development” and “Gene Expression in β-cells”, while “Signaling by Receptor Tyrosine Kinases” was unexpectedly up-regulated (Figure 3F).

**Figure 3.**
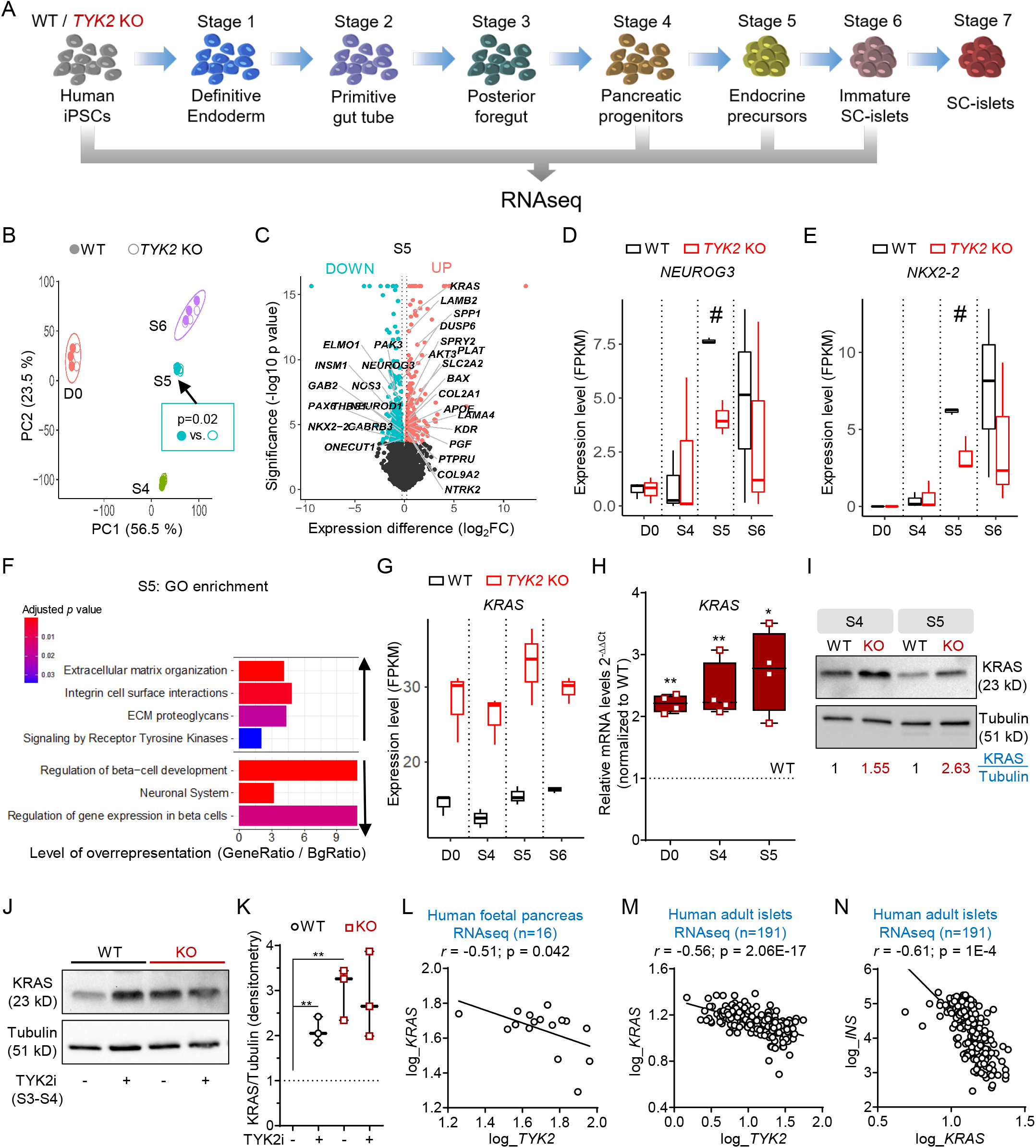
*TYK2* negatively regulates *KRAS* expression. **(A)** Schematic for the bulk RNAseq analysis of differentiating iPSCs. Samples were collected from 3 separate experiments with WT and *TYK2* KO cells (n = 3). **(B)** Principal component (PC) analysis of bulk RNA-seq comprising all samples. Filled circles indicate WT cells while empty circles indicate KO cells. **(C)** Volcano plot for Stage 5 differentially expressed genes between WT and *TYK2* KO, significantly downregulated genes are indicated in iris blue and upregulated genes in soft red. Selected genes of interest are highlighted. **(D-E)** Boxplot showing FPKM values for the expression of *NEUROG3* and *NKX2-2* at indicated Stages of differentiation. **(F)** Selected up- and down -regulated Reactome enrichment pathways in Stage 5 *TYK2* KO cells compared to WT. **(G)** Boxplot showing FPKM values for the expression of *KRAS.* **(H)** Relative transcript levels of *KRAS* determined with qRT-PCR in hiPSCs, Stage 4 and Stage 5 cells (n = 4). Box and whiskers plots showing min to max with all the points. **(I)** Immunoblot analysis of KRAS protein at Stage 4 and Stage 5. Normalized (with tubulin) densitometric values indicated in the panel (n = 3). **(J)** Immunoblot analysis of KRAS protein at Stage 4 in WT and *TYK2* KO cells following TYK2i (BMS986165) treatment during Stage 3 to 4. **(K)** Densitometric analysis of panel J (n = 3). **(L)** Correlation of *TYK2* and *NEUROG3* expression in RNA-seq of human fetal pancreas samples. Pearson’s correlation after log normalization of counts (n = 16). **(M)** Correlation of *KRAS* and *TYK2* expression; and **(N)** *KRAS* and *INS* expression in human islet RNA-seq samples (n = 191). Pearson correlations *r* and significance levels *p* are also indicated in the panels. Two-tailed unpaired t-test was applied to determine the significance levels unless otherwise mentioned. *p < 0.05; **p < 0.01.

There were only 16 consistently up-regulated and 3 down-regulated genes throughout all stages of differentiation. Strikingly, we observed that the most significantly up-regulated gene at all stages was *KRAS* (Figure 3G). This observation was confirmed at gene (qRT-PCR) and protein (WB) levels (Figure 3, H and I). Similarly, we observed a significant increase of KRAS protein upon TYK2i treatment in WT, but not KO cells, indicating a negative correlation between TYK2 and KRAS (Figure 3, J and K). The RNAseq data from human fetal pancreata (r=- 0.51; *p*=0.042) and adult islets (r=- 0.56; *p*=2.06E-17) also showed a significant negative correlation between *TYK2* and *KRAS* expression levels (Figure 3, L and M). Moreover, a strong negative correlation between *KRAS* and *INS* (r =-0.61; *p*=1x10^-4^) was also found in the islet dataset (Figure 3N).

Collectively, these findings are in agreement with earlier studies showing that KRAS antagonizes endocrine neogenesis in the developing pancreas through inhibition of NEUROG3 expression(19) and suggest that TYK2 loss or inhibition compromises EP formation by upregulating KRAS.

### Increased *KRAS* leads to shortening of G_1_ phase and inhibition of *NEUROG3* expression in *TYK2* KO EP cells

To understand the impact of *TYK2* KO at the level of specific pancreatic cell subtypes, we investigated the *TYK2* KO phenotype via single-cell (sc) RNA-seq (Figure 4A). We generated a dataset of 12545 WT and 13856 KO cells for EP (S5) and SC-islets (S6), and analysed it using the Seurat pipeline(20). After quality filtering, we annotated the clusters according to the expression of known cell-type specific pancreatic markers (Figure 4, B and C and Supplemental Figure 4, A and B). Next, we classified the clusters broadly in four groups: PP (*SOX9*^++^*ONECUT1*^++^*TCF7L2*^++^), EP (*NEUROG3*^++^), α-like cells (*GCG*^++^*ARX*^++^) and β-like cells (*INS*^++^*NKX6-1*^++^). We observed a marked difference in the normalized distribution of cells in the clusters. WT S6 contained 62.2% β-like cells compared to 37.7% in KO, whereas S6 KO had still 64.4% PP-like cells compared to 35.5% in WT (Figure 4, D and E). GSEA (Reactome) analysis for PP and EP clusters at S5 confirmed the downregulation of the gene set for “Regulation of β-cell Development”, whereas gene sets for “M Phase” and “Cell Cycle” were upregulated in KO (Figure 4F). Furthermore, we confirmed the reduced expression of *NEUROG3* and its downstream target *NKX2-2* in KO EP cells, in line with our previous observations (Figure 4G).

**Figure 4.**
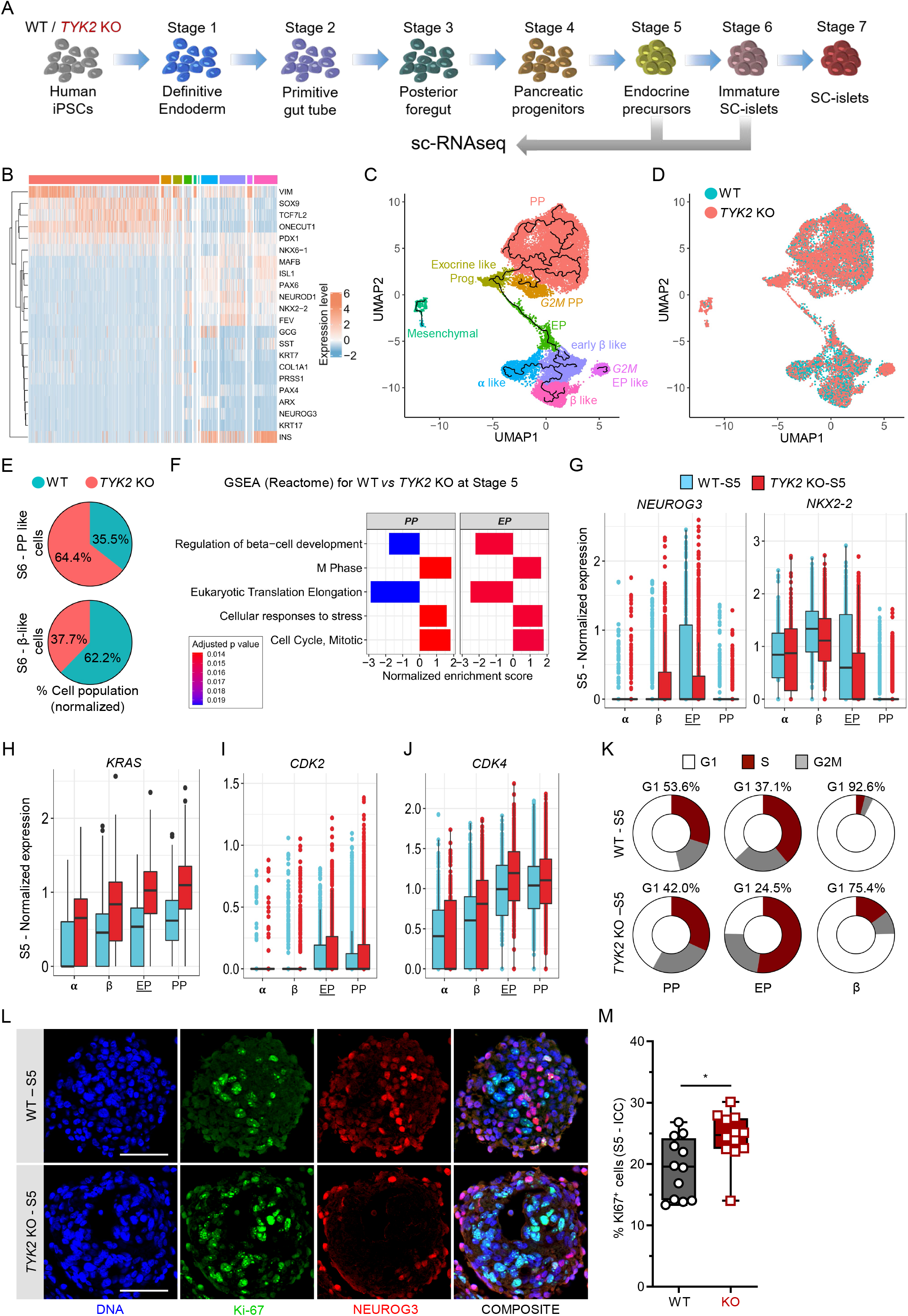
Single-cell transcriptomic analysis of endocrine differentiation. **(A)** Schematic for the scRNA-seq performed at Stage 5 (3995 WT and 5025 KO -cells) and Stage 6 (3348 WT and 4550 KO cells) samples. **(B)** Heatmap for the selected genes associated with pancreatic differentiation. **(C)** Various noted clusters are indicated in different color codes and presented as Uniform Manifold Approximation and projection (UMAP) with pseudotime trajectories. **(D)** Superimposed UMAP of WT in iris blue and *TYK2* KO in soft red. **(E)** Pie charts showing the normalized percentages of pancreatic progenitors (top) and β-like cells (bottom) in WT (iris blue) and *TYK2* KO (soft red) samples at the end of Stage 6. **(F)** scRNA-seq of Stage 5 WT (n = 3995) and KO (n = 5025) cells shown for selected enriched pathways in PP and Endocrine Precursors (EP) like clusters using Gene set enrichment analysis (GSEA, Reactome). **(G)** Boxplots for the relative expression of *NEUROG3* and *NKX2-2*; **(H)** *KRAS* **(I)** *CDK2* **(j)** *CDK4* in indicated α-like cells (α), β-like cells (β), EP and PP like clusters. **(K)** Donut chart showing the percentage of cells at various cell cycle phases with Seurat (CellCycleScoring) pipeline in PP, EP, and β-like cells cluster. **(L)** Representative image showing immunocytochemistry for NEUROG3 and Ki-67 co-expression at Stage 5. **(M)** Quantification of the data in panel L presented with box and whiskers plots showing min to max and all data points. Two-tailed unpaired t-test was performed to determine the significance levels. *p < 0.05. Scale bar = 100μm.

scRNA-seq revealed that the expression of *KRAS* was highest in the PP- and EP-like *TYK2* KO clusters (Figure 4H). *KRAS* has been recently shown to accelerate the G_1_/S transition of the cell cycle, leading to the shortening of G_1_ length (21). Lengthening of the G_1_ phase in the PP cells is a key to the proper augmentation of *NEUROG3* and its downstream targets(22). We observed a higher expression of cell division protein kinase 2 (*CDK2)* and *CDK4* in the *TYK2* KO S5 PP and EP cells (Figure 4, I and J). We then used the expression level of cell cycle markers to estimate the fraction of dividing cells in each clusters with Seurat (CellCycleScoring) (20). Notably, 53.6% and 37.1% cells of S5 WT PP and EP clusters, respectively, were in G_1_ phase compared to 42% and 24.5% in KO (Figure 4K).

Collectively, these data indicate that elevated KRAS drives the G_1_/S transition faster in the *TYK2* KO PP and EP cells, resulting in a G_1_ shortening that interferes with the proper induction of *NEUROG3* expression. Accordingly, we observed a significantly (*p*=0.013) higher Ki-67 nuclear staining in the KO S5 clusters (Figure 4, L and M).

### TYK2 regulates the IFNα response in SC-islets

Several studies reported the prominent role of IFNα in the induction of autoimmunity in T1D(23). Since TYK2 is important for the IFN-I signaling mediated through IFNAR1, we took advantage of our *TYK2* KO model to understand its role during the dialogue between IFNα and the developing pancreatic islet cells. The phosphorylation of STAT1/2 was completely abrogated in the S6 KO SC-islets in response to IFNα but not IFNγ, whereas STAT3 responses were only partially affected (Figure 5A).

**Figure 5.**
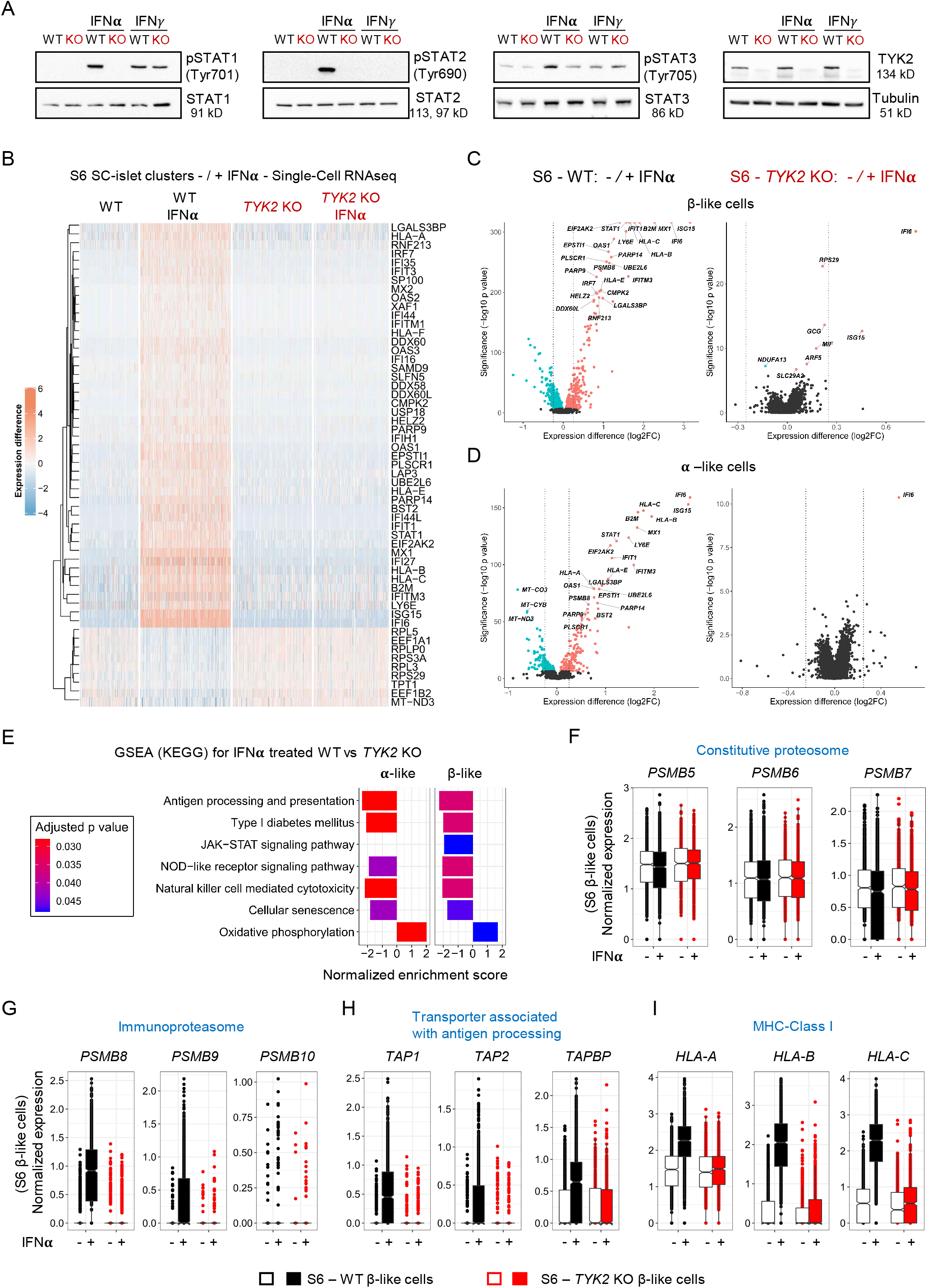
TYK2 regulates the IFNα responses in SC-islets. **(A)** Immunoblot based analysis for the phosphorylation of STAT1, STAT2 and STAT3 in Stage 6 SC-islets in response to either IFNα or IFNγ treatment (n=3). **(B)** Heatmap showing the differential expressed genes in the WT (3348 cells), WT+IFNα (5202 cells), KO (4550 cells) and KO+IFNα (4281 cells) samples. Single-cell transcriptomics performed following 24h IFNα (100 ng/ml) treatment on the WT and *TYK2* KO S6 SC-islets. **(C)** Volcano plot showing the significant upregulated (soft Red), downregulated (iris blue) and non-significant (black) genes in response to IFNα treatment in β-like cells and **(D)** for α-like cells. **(E)** GSEA (KEGG) based selected enriched gene sets shown in α- and β- like cells in response to IFNα treatment. **(F)** Condensed Boxplot showing the normalized expression of constitutive proteasome genes in β-like cells. **(G)** Similar presentation for the immunoproteasome genes and **(H)** Transporter associated with antigen processing genes and **(I)** MHC Class Ι genes.

Next, we performed scRNA-seq on S6 WT (5202 cells) and KO (4281 cells) islet cells treated with IFNα. Notably, the global transcriptomic changes following IFNα treatment in WT individual islet cells were in good agreement with our previously reported human islet dataset under similar treatment(5) (Supplemental Figure 5A). IFNα induced the up-regulation of previously described IFN-stimulated genes (ISGs) in the WT cells. However, this response was absent in *TYK2* KO cells, shown by a heatmap for the top 55 (46 up and 9 down) differentially expressed genes in all clusters (Figure 5B and Supplemental Figure 5B). The IFNα response was similarly completely inhibited in both β- and α-like cells (Figure 5, C and D). Furthermore, genes important for antiviral responses to IFNs like *STAT1* and antiviral MX dynamin-like GTPase 1 (*MX1*) were strongly induced in WT but remained unchanged in the *TYK2* KO cells (Supplemental Figure 6A). Given that MHC Class Ι up-regulation is one of the most important hallmarks in diabetic β-cells(24), we observed that upon IFNα treatment, *HLA-A*, *HLA-B*, *HLA-C* and *HLA-E* genes were strongly up-regulated in WT cells, but unchanged in *TYK2* KO cells (Supplemental Figure 6B). In agreement with the experimental data, a strong positive correlation between *TYK2* and *HLA-A* (r=0.52; *p*=6.53x10^-15^); *HLA-B* (r=0.42; *p*=1.06x10^-9^); *HLA-C* (r=0.60; *p*=1.55x10^-20^); *HLA-E* (r=0.51; *p*=3.65x10^-14^) was observed in the human islet RNA-seq dataset (n=191)(17) (Supplemental Figure 6C).

In line with the other findings described above, GSEA (KEGG) analysis highlighted lower expression of antigen processing and presentation gene sets in *TYK2* KO β- and α-like cells following IFNα compared to WT (Figure 5E). We then analysed the expression of genes associated with antigen processing following IFNα treatment. First, we observed that constitutive proteasome genes (*PSMB5*, *PSMB6* and *PSMB7*) remained unchanged in both WT and KO while IFNα-upregulated genes encoding the immunoproteasome (*PSMB8, PSMB9* and *PSMB10*). Second, transporters associated with antigen processing (*TAP1, TAP2* and *TAPBP*) and MHC Class Ι (*HLA-A, HLA-B and HLA-C*) were strongly augmented in the WT β-like cells but remained unchanged in the KO(25) (Figure 5, F-I).

Collectively, these data demonstrate that inhibition of TYK2 signaling in β- and α-like cells inhibited both antigen processing and presentation following IFNα treatment. Because of the partially compromised endocrine differentiation of *TYK2* KO SC-islets, these findings could also be related with a lower expression of T1D associated β-cell autoantigens, notably insulin. We therefore compared the expression of previously described T1D autoantigens between WT and *TYK2* KO β-like cells(26). A significantly lower expression was found only for *GAD1* (in the presence of IFNα) and *GAD2* in *TYK2* KO β-like cells (Supplemental Figure 6, D and E).

### TYK2 modulates MHC Class Ι presentation and T-cell-mediated cytotoxicity in SC-islets

Next, we examined the expression of MHC Class Ι protein on the surface of maturing SC-islets following IFNα treatment. We observed a strong up-regulation on the WT SC-islets, whereas this response was absent in *TYK2* KO or upon TYK2i co-treatment (Figure 6, A and B). Similarly, flow cytometry confirmed the overall 83.1% MHC Class Ι increase in WT SC-islets compared to 0.69% in *TYK2* KO (Figure 6, C-E). Notably, *TYK2* KO SC-islets also had 26.9% INS^+^ cells compared to 41.8% in WT SC-islets.

**Figure 6.**
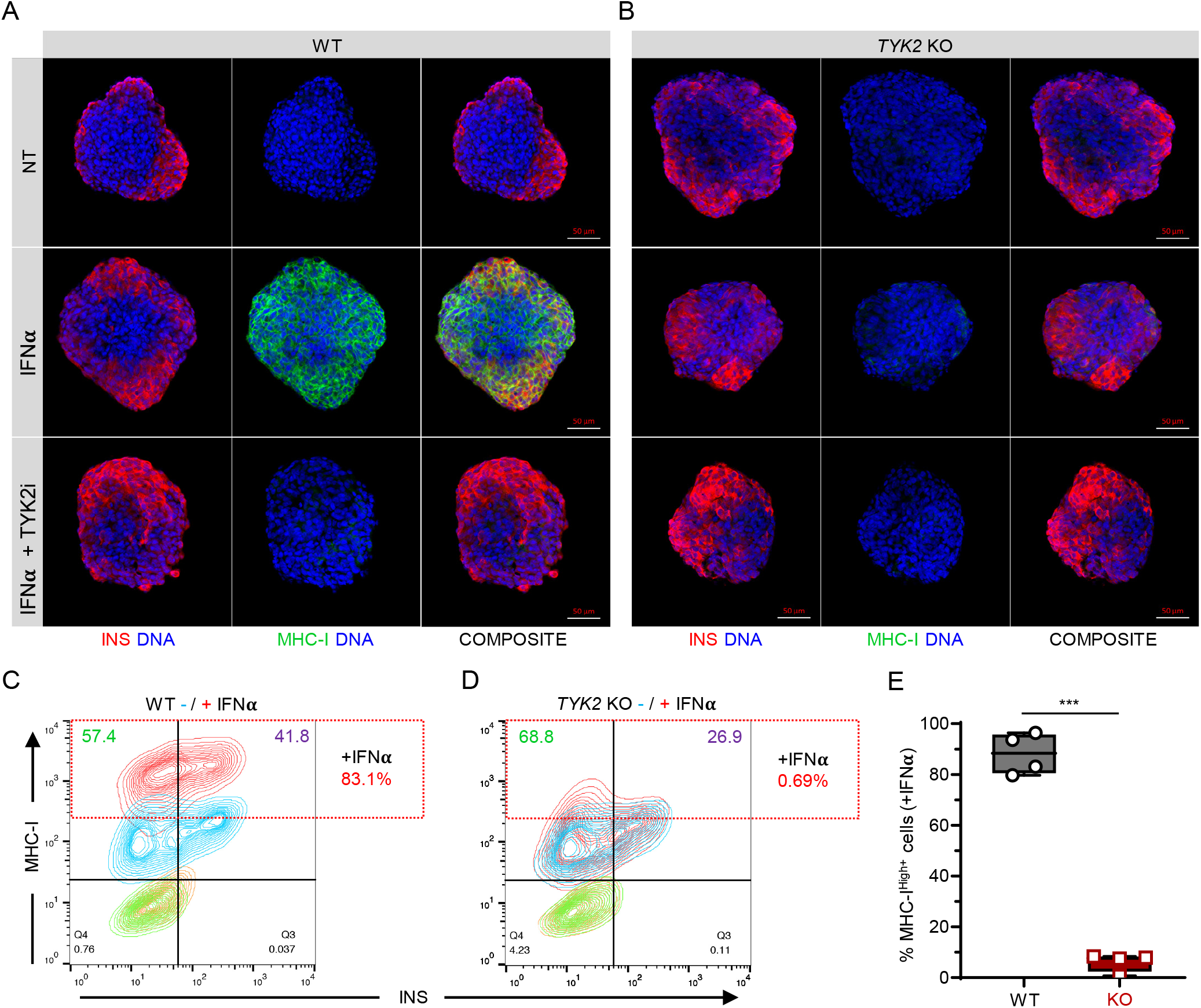
TYK2 modulates MHC Class Ι presentation in SC-islets. **(A-B)** Representative images of S7 SC-islets showing the immunocytochemistry for insulin (red) and MHC Class Ι (green) expression following 24 h treatment with IFNα alone or in combination with TYK2i. Panel A, WT cells; panel B, *TYK2* KO cells. **(C-D)** Representative contour plots of flow cytometry showing the co-expression of insulin and MHC Class Ι following 24h IFNα treatment. Panel C, WT cells; panel D, *TYK2* KO cells. **(E)** Quantification of the data in C-D. Two-tailed unpaired t-test were performed to determine the significance levels. Box and whiskers plot showing min to max with all the points. ***p < 0.001. Scale bar = 50 μm.

To investigate whether inhibiting IFNα-induced MHC Class Ι up-regulation by TYK2i could reduce the vulnerability of WT SC-islets to T-cell-mediated cytotoxicity, we designed a T-cell co-culture experiment (Figure 7A). To exclude the potential confounding effect of β-cell antigen downregulation upon TYK2 loss, we used a flu peptide-reactive CD8^+^ T-cell line. Variable T-cell numbers were cultured for 6 h with a fixed mixture of peptide-pulsed, CFSE- labeled SC-islets and unpulsed, CTV-labeled SC-islets, which were pre-treated with IFNα alone or in combination with TYK2i or left untreated. The ratio of surviving SC-islets thus provided a readout of peptide-specific lysis (Figure 7B). IFNα treatment led to a significant increase in SC-islet lysis, which was inhibited in the presence of TYK2i (Figure 7C). These cytotoxic outcomes were paralleled by MHC Class Ι expression (Figure 7, D and E). With increasing T-cell numbers, SC-islets that survived cytotoxicity were those with lower MHC Class Ι expression (Figure 7E). Although the inhibition of MHC Class Ι up-regulation by TYK2i was complete, cytotoxicity was only partially inhibited. In line with the known effect of IFNα on programmed death ligand-1 (PD-L1) up-regulation(27) TYK2i treatment also partially inhibited PD-L1 up-regulation (Figure 7, D and F). PD-L1^high^ SC-islets were those surviving cytotoxicity with increasing T-cell numbers up to 5:1 effector-to-target ratios (Figure 7F). The concomitant inhibitory effect of TYK2i on MHC Class Ι and PD-L1 up-regulation may explain the incomplete inhibition of T-cell mediated cytotoxicity.

**Figure 7.**
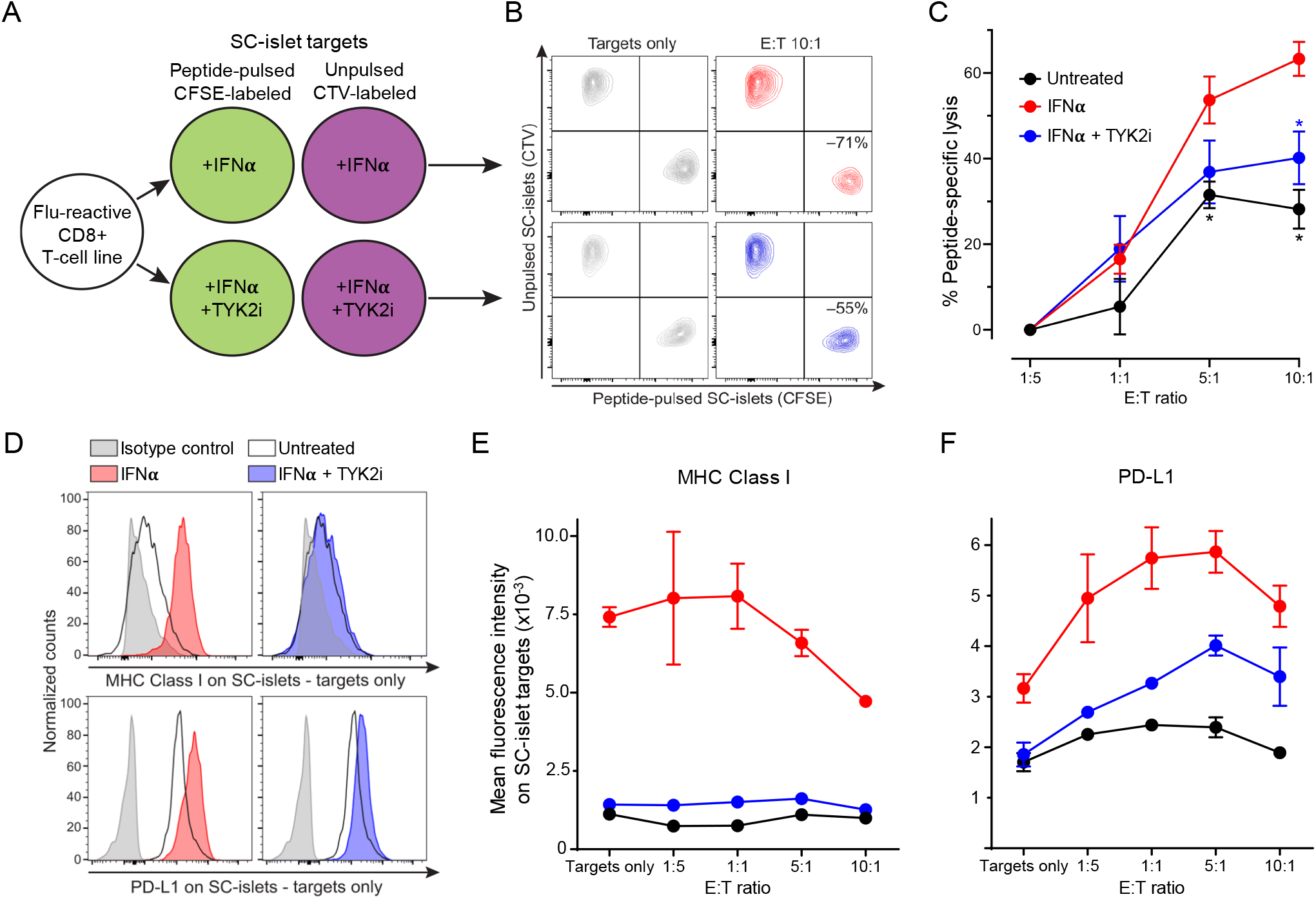
TYK2 inhibition modulates T-cell-mediated cytotoxicity in SC-islets. **(A)** Schematic of T-cell/SC-islet co-culture experiments. Increasing numbers of flu peptide-reactive CD8^+^ T cells were incubated for 6 h with a fixed mixture of peptide-pulsed, CFSE- labeled SC-islets and unpulsed, CellTrace Violet (CTV)-labeled SC-islets, which were preliminary treated for 24 h with IFNα alone or in combination with TYK2i or left untreated. The ratio of surviving (Live/Dead-negative) pulsed CFSE^+^ *vs.* unpulsed CTV^+^ SC-islets thus provided a readout of peptide-specific lysis. **(B)** Representative flow cytometry dot plots of IFNα-treated (top) and IFNα/TYK2i-treated SC-islets (bottom), cultured alone (left) or with T cells at an effector-to-target (E:T) ratio of 10:1 (right). Each population is gated on Live/Dead-negative events. The percent peptide-specific lysis in the presence of T cells (i.e., the ratio of CFSE^+^/CTV^+^ live cells normalized to the ratio in the absence of T cells) is indicated. **(C)** Peptide-specific lysis for the indicated conditions at different E:T ratios. Data are normalized means ± SEM of two experiments performed in triplicate: \**p*<0.05 by Mann-Whitney U test vs IFNα. **(D)** Representative flow cytometry histograms of MHC Class Ι (top) and PD-L1 expression (bottom) in SC-islet targets treated with IFNα (left) or IFNα/TYK2i (right). **(E-F)** Mean fluorescence intensity of **(E)** MHC Class Ι and **(F)** PD-L1 expression in SC-islets at different E:T ratios. Data are means ± SEM of duplicate wells and one representative experiment out of two performed is shown.

Collectively, these results demonstrate that TYK2 inhibition prevents IFNα-induced MHC Class Ι up-regulation on SC-islets and, despite some concomitant inhibition of PD-L1 up- regulation, significantly decreases the ensuing T-cell-mediated cytotoxicity.

### Replication and analysis of loss-of-function *TYK2* SNP (rs34536443) in R5 FinnGen cohort

Loss-of-function SNPs of *TYK2* (e.g., rs34536443; TYK2^P1104A^) have been reported to be protective against several autoimmune diseases including T1D(9, 28). The TYK2i BMS- 986165 used in the present study mimics the effect of the TYK2^P1104A^ coding variant(18). We replicated the phenome-wide association analysis for this SNP in the R5 dataset of FinnGen project (r5.finngen.fi), which compiled 2803 clinical endpoints obtained from electronic health record data of 218,792 Finnish individuals. We found that rs34536443 provided protection from several autoimmune/auto-inflammatory diseases in the Finnish population, including Crohn’s disease, seropositive rheumatoid arthritis, sarcoidosis, autoimmune hypothyroidism and T1D (5,928 cases and 183,185 controls; p<1.16x10^-5^) (Supplemental Figure 7).

## Discussion

Here we demonstrate a previously unknown role of the T1D candidate gene *TYK2* in pancreatic endocrine cell development. In the absence of TYK2, KRAS expression is upregulated, which results in improper induction of the pro-endocrine transcription factor NEUROG3. However, at the mature stage SC-islets expand and function normally in the absence of TYK2 (Figure 2I and Supplemental Figure 2, G and H). Importantly, TYK2 was found to be essential for the IFN-Ι responsiveness of SC-islets, supporting the therapeutic rationale of TYK2 inhibition to halt T1D progression.

Directed differentiation of hPSCs into pancreatic islet cells provides a controlled experimental system to study the role of diabetes-associated genes in pancreas development(29, 30). We and others have previously shown that STAT3 activation regulates the NEUROG3 mediated pancreatic β-cell differentiation(12, 31). In the present study, we show that TYK2 is a major STAT activator that also has a regulatory role on NEUROG3 expression in pancreatic progenitors. Interestingly, proinflammatory cytokines (IL1β, TNFα and IFNγ) induce endocrine differentiation in pancreatic ductal cells through STAT3 dependent NEUROG3 expression(32).

Our results revealed an unexpected and previously unknown negative regulation between *TYK2* and *KRAS* (Figure 3, J-M). Regulation of KRAS expression by TYK2 in KO cells is noteworthy, since JAK/STAT signaling is known to be important for receptor tyrosine kinase (RTK) and mitogen-activated protein kinase (MAPK) signaling(33). In agreement with this, Reactome enrichment analysis of S5 bulk RNA-seq revealed a significant up-regulation of RTK pathways in the *TYK2* KO cells. It has been shown that KRAS accelerates the G_1_/S progression, and lengthening of the G_1_ phase is a prerequisite for the complete induction of NEUROG3 expression before cell-cycle exit of the endocrine progenitors(19, 22). Thus, an elevated KRAS containing *TYK2* KO progenitors have a shorter G_1_ phase which results into a compromised NEUROG3 augmentation (Figure 4). Taken together, we show that regulation of the JAK/STAT pathway in pancreatic progenitors by TYK2 is essential for the cell-cycle control required for the progression of endocrine differentiation.

Conspicuously, rare loss-of-function *TYK2* promoter mutations (Clin Var, 440728) in the Japanese population predispose to autoantibody-negative T1D and T2D (8). This association has been thought to be linked with increased susceptibility to viral infection(34). However, based on our results, the partial loss of TYK2 expression could lead to a lower β-cell mass as a contributing factor to the increased diabetes risk.

Notably, a germline loss-of-function *TYK2* mutation (rs34536443; TYK2^P1104A^) has been reported and hereby confirmed in the Finnish population to be protective against several autoimmune diseases including T1D(9, 28) (Supplemental Figure 7). This highly protective effect against autoimmunity is associated with the dampening of IFN-Ι, IL12 and IL23 signaling(28). Of note, IFN-Ι responses are important contributors to T1D etiology and inhibition of these responses in early disease has been suggested as a potential intervention(6, 35). *TYK2*-silenced adult β-cells (i.e. a 50% inhibition induced by siRNA or TYK2i) displayed limited IFNα pathway activation, but preserved antiviral responses and β-cell function(10). Our experiments showed that loss of TYK2 abolished the up-regulation of ISGs including MHC Class I, equivalently in β-cells, α-cells and their EPs (Figure 5, C and D and Supplemental Figure 6, A and B). Additionally, the IFNα-induced up-regulation of antigen processing genes, i.e. immunoproteasome (*PSMB8*, *PSMB9*) and peptide transporters (*TAP1*, *TAP2* and *TAPBP*) were lost, suggesting impairment of the processing and presentation of immunogenic peptides in the *TYK2* KO cells(25). It is thus possible that these effects may synergize to inhibit autoimmune responses *in vivo* to a larger extent than what we observed *in vitro* using peptide-reactive CD8^+^ T cells against peptide-pulsed SC-islets (Figure 7). Despite the higher cytotoxic potency of these peptide-driven responses compared to the autoimmune responses against naturally processed and presented β-cell peptides, and the concomitant inhibition of IFNα-induced PD-L1 up-regulation, TYK2i was still effective at limiting SC-islet vulnerability to T-cells. Careful TYK2i titration, or concomitant targeting of other downstream mediators of IFNα signaling, may allow to dissociate its inhibitory effects on MHC Class Ι and PD-L1.

A recent report suggests that iPSC-α cells are selectively protected from T-cell mediated destruction compared to iPSC-β cells following co-culture with autologous PBMCs, albeit they were immunogenic in an allogenic setting(36). Importantly, our scRNA-seq dataset revealed a similar *TYK2* dependent up-regulation of ISGs in α-cell- and β-cell-like populations, indicating a similar regulatory mechanism. The present model may thus prove useful to understand how α-cells are selectively protected from T-cell mediated autoimmunity despite having similar IFNα responses.

In summary, using *TYK2* KO SC-islet models, we deciphered the dual role of the candidate gene *TYK2* in pancreatic β-cells. First, depletion of TYK2 during early islet development affected the endocrine commitment, while it did not affect the functionality of mature beta cells. Second, TYK2 inhibition in mature islet cells effectively inhibited the IFNα induced up- regulation of the antigen processing and presentation machinery, which reduced vulnerability to T-cell cytotoxicity. Inhibiting TYK2 signaling was sufficient to protect SC-islets from IFNα responses and T-cell cytotoxicity. Importantly, the TYK2i BMS-986165 used in this study is already into phase 2 and 3 trials for psoriasis(18, 37) and may thus provide an attractive candidate for T1D interventions.

## Methods

### hPSCs Cell lines and Genome editing

hiPSCs line HEL46.11 (derived from human neonatal foreskin fibroblast)(38) and Human embryonic stem cell line H1 (WA01, WiCell) were used in the study. HEL46.11 hiPSCs were used to generate the *TYK2* knockout lines used in this study. The hPSCs were cultured on Matrigel (Corning, #354277) coated plates in Essential 8 (E8) medium (Thermo Fisher, A1517001) and passaged with 5 mM EDTA (Thermo Fisher, #15575-038) in PBS. The cell lines were routinely tested for the mycoplasma contamination.

For knocking out *TYK2* in HEL46.11 iPSCs, the ATG starting codon-containing third exon was deleted using two CRISPR/Cas9 guides that were designed with Benchling (Biology Software, 2017) (**G1** AAGAGCTAACAGGGGTCTCT and **G2** GTCTGGGGCGTTGGCACCAT). Two million iPSCs were electroporated with 6 μg of CAG- Cas9-T2A-EGFP-ires-Puro (deposited in Addgene, plasmid no. 78311, together with detailed protocols for its use), 500 ng of gRNA1-PCR *TYK2* and 500 ng of gRNA2-PCR *TYK2* products, using Neon Transfection system (Thermo Fisher Scientific; 1100 V; 20 ms; two pulses). Single cells were sorted, expanded, and screened for approximately 300 bp deletion of exon 3. Positive clones were validated by Sanger sequencing. The top three CRISPR off-target hits were sequenced and did not have any indels. Primers sequences used in the study to for genome editing is described in supplementary table.

### *In vitro* hPSCs culture and their pancreatic lineage differentiation

hPSCs were differentiated towards pancreatic lineage to generate SC-islets using our recently published detailed protocol(11) with minor modifications. Briefly, near 80%confluent plates of stem cells were dissociated using EDTA and seeded on new Matrigel coated plates in E8 medium supplemented with 10 μM Rho- Associated kinase inhibitor (ROCKi, Y-27632, Selleckchem S1049) at a density of approximately 0.22 million cells/cm^2^. The differentiation was started following 24 h of seeding and proceeded through a 7-stage differentiation protocol (stages 1 – 4 in adherent culture, stage 5 in AggreWell (Stemcell Technologies, #34421), and stages 6 and 7 in suspension culture).

The detailed differentiation protocol and the stage specific complete media formulations are described below. The details of the reagents used in the differentiation protocol described in the supplementary table.

| Stage | Reagent | Medium |
| --- | --- | --- |
| Stage 1<br>3 days | d0: 100 ng/ml human Activin A, 3 μM CHIR<br>d1: 100 ng/ml human Activin A, 0.3 μM CHIR<br>d2: 100 ng/ml human Activin A | Basal 1: MCDB131 (10372-019, Life Technologies), 2 mM Glutamax, 1.5 g/L NaHCO<sub>3</sub> (Sigma-Aldrich), 0.5% BSA fraction V Fatty acid free (Sigma-Aldrich), 10 mM glucose (Sigma-Aldrich) |
| Stage 2<br>3 days | 0.25 mM Ascorbic acid, 50 ng/ml FGF7 |  |
| Stage 3<br>2 days | 0.25 mM Ascorbic acid, 50 ng/ml FGF7, 0.25 μM SANT1, 1 μM Retinoic Acid, 100 nM LDN, 200 nM and TPB | Basal 2: MCDB131, 2 mM Glutamax (35050038, Life Technologies), 2.5 g/L NaHCO<sub>3</sub>, 2% BSA fV, 10 mM glucose, 1:200 ITS-X (51500-056, Life Technologies) |
| Stage 4<br>4 days | 0.25 mM Ascorbic acid, 50 ng/ml FGF7, 0.25 μM SANT1, 0.1 μM Retinoic Acid, 200 nM LDN, 100 ng/ml EGF and 10 mM Nicotinamide | |
| Stage 5<br>4 days | 0.25 μM SANT1, 0.05 μM Retinoic acid, 100 nM LDN, 10 μM ALK5inhII, 1 μM GC1, 20 ng/mL Betacellulin and 100 nM GSiXX | Basal 3: MCDB131, 2 mM Glutamax, 1.5 g/L NaHCO<sub>3</sub>, 2% BSA fV, 20 mM glucose, 1:200 ITS-X, 10 μg/mL Heparin (H3149, Sigma-Aldrich), 10 μM Zinc Sulfate (Z0251, Sigma-Aldrich) |
| Stage 6<br>8 days | 100 nM LDN, 10 μM ALK5inhII, 1 μM GC1 and 100 nM GSiXX | |
| Stage 7<br>21 days | 1 mM N-Acetylcysteine, 10 nM Tri-iodothyronine (T3) and 0.5 μM Aurora kinase inhibitor ZM447439 | CMRL-modified: CMRL 1066 (15-110-CVR, Corning), 2% BSA fV, 2 mM Glutamax, 1:200 ITS-X, Heparin 10 μg/ml, 10 μM Zinc Sulfate, 5 mM Sodium Pyruvate (Lonza), 1:2000 chemically defined lipid concentrate (11905-031, Invitrogen), 1:2000 medium trace elements A (25-021-CI, Cellgro), 1:2000 medium trace elements B (99-175-CI, Cellgro) |

### Human pancreatic islets RNAseq for correlation studies

Data from human pancreatic islets were processed as described(17). Briefly, human islets (n = 191) were obtained through the EXODIAB network from the Nordic Network for Clinical Islet Transplantation. 28 T2D donors had a clinical diagnosis of T2D. RNA was extracted using miRNeasy (Qiagen) or the AllPrep DNA/RNA (Qiagen) mini kits miRNeasy (Qiagen) or the AllPrep DNA/RNA (Qiagen) mini kits. Library preparation of high-quality RNA (RIN >8) was performed using the TruSeq RNA sample preparation kit (Illumina). Raw sequencing data was aligned to human genome B38. Gene length adjusted counts (FPKMs) was used to calculate gene-gene correlations using the Pearson method.

### Human fetal pancreas

RNA was extracted from tissue biopsies from terminated foetuses (7–14 gestational weeks) using the TRIZOL method (n = 16). RNA libraries were then constructed using the TruSeq RNA library preparation kit (Illumina). RNA sequencing was performed on a HiSeq 2000 / Nextseq (Illumina). Paired-end 101 bp-long reads were aligned to the Reference Human Genome Build 37 using STAR and counts were generated as described(39). Batch corrections were performed using COMBAT. Expression levels between fetal and adult (from GTEX) pancreas were compared in EdgeR. Gene expression was related to days post-conception using Pearson correlation. Gene-gene correlations were performed using Pearson correlation after log normalization of counts.

### Ultra-deep bulk RNAseq analysis

We performed ultra-deep RNAseq analysis for the HEL46.11 derived stage 1 (DE), stage 4 (PP) and stage 7 (SC-islets) samples shown in Figure 1B and C. The differentiation, RNA isolation and library preparation was performed as described previously(12). Sequencing was performed using Illumina HiSeq 2500 (with chemistry v4) at Eurofins Genomics.

The read pairs were mapped to the human reference genome (GRCh38) with STAR aligner(40). Gene expression was counted from read pairs mapping to exons using featureCounts in Rsubread(41). Duplicates, chimeric and multimapping reads were excluded, as well as reads with low mapping score (MAPQ <10). The read count data was analysed with DESeq2(42). We analysed the effect of differentiation as a function of time, as well as pairwise comparisons between the different developmental stages (DE - PP-SC-islets). We removed genes with low expression levels from the analysis (<50 reads across all samples). PCA was calculated with prcomp using normalized counts that were scaled using the voom-function from limma(43).

For WT and *TYK2* KO genotype samples, RNAseq (Illumina HiSeq 2500; v4)) was performed at stage 0 (iPSCs), stage 4 (PP), stage 5 (EP) and stage 6 (EC) as presented in Supplemental Figure 1 and Figure 3. The raw data was filtered with Cutadapt to remove adapter sequences, ambiguous (N) and low-quality bases (Phred score <25). We also excluded read pairs that were too short (<25bp) after trimming. The filtered read pair were mapped to the human reference genome (GRCh38) with STAR aligner(40) and processed as described above. We analysed the effect of *TYK2* knockout separately for each developmental stage (iPSCs, stage 4, stage 5, stage 6). For false discovery rate (FDR) estimation we used Fdrtool(44). The differentially expressed genes (FDR<0.01) were analyzed for enrichment separately for the up- and down-regulated genes using ClusterProfiler(45) against the Reactome pathways(46).

### Single cell RNA sequencing sample preparation and analysis

WT and *TYK2* KO genotype stage 5 and stage 6 (either untreated or with 100 ng/ml for 24 h IFNα treatment) samples were prepared for the single cell RNAseq analysis as we described previously(11).

Single cell gene expression profiles were generated with 10x Genomics Chromium Single Cell 3’RNAseq platform using the Chromium Next GEM Single Cell 3’ Gene Expression (version 3.1 chemistry). Raw data (fastq) processing was performed with 10x Genomics Cell Ranger (v3.1) pipeline. The reads were mapped to the human reference genome (GRCh38.98). The filtered counts were analysed with Seurat(20). The counts were normalized, scaled, and analysed for PCA with default methods. The variable genes (top 1000) were identified separately for each sample and combined during the analysis (for a total of 1199 variable genes). To reduce biases among datasets we used Harmony(47) on the first 50 PCs with sample as the covariable (with theta = 2, nclust = 50, max.iter.cluster = 40, max.iter.harmony = 10). The integrated (harmonized) PCs were used to build the UMAP, find the neighboring cells (using Shared Nearest Neighbor), and identify cell clusters using default Seurat methods. To reduce background RNA contamination from disrupted cells we used SoupX(48) with clusters identified with Seurat, and known cell type specific marker genes (GCG, TTR, INS, IAPP, SST, GHRL, PPY, COL3A1, CPA1, CLPS, REG1A, CELA3A, CTRB1, CTRB2, PRSS2, CPA2, KRT19, VTCN1) to estimate the level of contamination. The Seurat analysis was then repeated with the adjusted counts with the following modifications. Cells with less than 5000 UMI counts or 1700 expressed genes were excluded. We also removed cells with unusually high level of mitochondrial reads (>20% of counts). We assigned a cell cycle phase to all the cells using the default settings in CellCycleScoring-function and used this to remove biases due to cell cycle heterogeneity when scaling the data. During clustering the resolution was set to 0.2. Differentially expressed genes among clusters and sample types were identified with Wilcox-test using Seurat. The clusters were reordered by similarity and identified to cell types by the differentially expressed genes corresponding to known marker genes. The differentially expressed genes (FDR<0.01) were analyzed for enrichment separately for the up- and down- regulated genes using ClusterProfiler(45) against the KEGG. ClusterProfiler was also used for geneset enrichment analysis (GSEA) against the same database, using fold change to rank the genes.

### *In vivo* animal transplantation studies

Animal care and experiments were approved by National Animal Experiment Board in Finland (ESAVI/14852/2018). NOD-SCID-Gamma (NSG, 005557, Jackson Laboratory) mice were obtained from SCANBUR and housed at Biomedicum Helsinki animal facility, on a 12 h light/dark cycle and fed standard chow. Implantations were performed on 3- to 8-month-old mice as described previously(11). Briefly, stage 7- WT and *TYK2* KO SC-islets equivalent to approximately 2 million cells were loaded in PE-50 tubing and implanted under the kidney capsule. Mouse serum samples were collected monthly from the saphenous vein and stored at -80°C for human C-peptide analysis. Human-specific C-peptide was measured from plasma samples with the Ultrasensitive C-peptide ELISA kit (Mercodia, Uppsala, Sweden).

### Western blot

For protein extraction, cells were washed with ice-cold PBS and lysed with Cell lysis buffer (Cell Signaling Technologies #9803) for 10 min on ice. The cells were sonicated for 3 x 5 s on ice, centrifuged (1000 rcf for 10 min at 4°C) and the supernatant was stored at -80°C. The samples were run on Any kD Mini-PROTEAN TGX gel (Bio-Rad Laboratories) and then dry transferred onto a nitrocellulose membrane using the iBlot system (Invitrogen) as per manufacturer’s instructions. The membrane was then probed with the primary antibody overnight at 4°C, washed twice with Tris-buffered saline containing 0.05% Tween for 2 x 10 min, and incubated with the corresponding secondary antibody for 30 min at room temperature. Chemiluminescence detection was performed with Amersham ECL (RPN2235; Cytiva) and Bio-RAD Chemidoc XRS1 imaging system; Image Lab Software. The details of antibodies and their dilutions used in the study for WB are described in the supplementary table.

### Flow cytometry

For quantifying the definitive endoderm positive cells in stage 1, cytometry for CXCR4 was performed as previously described(12). For intracellular antigen cytometry of stage 7, cells were dissociated with TrypLE (Thermo Fisher Scientific) for 6 min at 37°C and resuspended in cold 5% FBS-containing PBS. Cells were fixed and permeabilized using Cytofix/Cytoperm (554714, BD Biosciences) as per manufacturer’s instructions. Primary or conjugated antibodies were incubated with the cells overnight at 4°C in Perm/Wash buffer (554714, BD Biosciences) containing 4% FBS and secondary antibodies for 45 min at RT. The cells were then analysed using FACSCalibur cytometer (BD Biosciences) and FlowJo software (Tree Star Inc.). The details of antibodies and their dilutions used in the study for flow cytometry are described in the supplementary table.

### mRNA extraction and qRT-PCR

Total RNA from hiPSC-derived cells were isolated using NucleoSpin Plus RNA kit (Macherey-Nagel). A total of 1.5 μg RNA was reversely transcribed using Moloney murine leukemia virus reverse transcriptase (M1701, Promega) for 90 min at 37°C. 50 ng cDNA was amplified using 5x HOT FIREPol EvaGreen qPCR Mix Plus no ROX (Solisbiodyne) in a 20 μl reaction. The reactions were pipetted using QIAgility (Qiagen) robot into 100 well disc run in Rotor-Gene Q. Relative quantification of gene expression was analysed using ΔΔCt method, with cyclophilin G (PPIG) as a reference gene. Reverse transcription without template was used as negative control and exogenous positive control was used as a calibrator. The qRT-PCR primers sequence will be made available from lead contact upon request.

### Immunocytochemistry and immunohistochemistry

For paraffin embedding, aggregates were fixed with 4% PFA at 4°C for 24h, following eosin staining, aggregates were embedded in 2% low-melting agarose (Fisher Bioreagents) PBS and transferred to paraffin blocks. WT or *TYK2* KO implanted grafts were retrieved at 2 or 3 months, dissected, and fixed with 4% PFA at RT for 48h, paraffin embedded, and cut into 5 μm sections using Leica microtome. For immunohistochemistry, slides were deparaffinized and antigens retrieved by boiling slides in 0.1 M citrate buffer (pH 6) using Decloaking chamber (Biocare Medical) at 95°C for 12 min. For whole mount or adherent cultures staining, cells were fixed in 4% PFA for 15 min at RT, permeabilized with 0.5% Triton X-100 in PBS for 15 min at RT, then blocked with Ultra-V (Thermo Fisher Scientific) for 10 min and incubated with primary antibodies overnight at 4°C and secondary antibodies for 1h at RT diluted in 0.1% Tween in PBS. Zeiss Axio Observer Z1 with Apotome was used to image the cells and further processed with ZEN-2 software. All stained samples were equally treated and imaged with the same microscope parameters. Image quantification was performed using CellProfiler software(49) and Fiji software(50). The details of antibodies and their dilutions used in the study for immuno-cytochemistry are described in the supplementary table.

### Static and dynamic glucose-stimulated insulin secretion

For the static assay, fifty aggregates were manually picked and preincubated in 2.8 mmol/L glucose Krebs buffer in a 12-well plate placed on a 95-rpm rotating platform for 90 min at 37°C. Aggregates were then washed with Krebs buffer and sequentially incubated in Krebs buffer containing 2.8 mmol/L glucose, 16.6 mmol/L glucose, and 2.8 mmol/L glucose plus 30 mmol/L KCL, for periods of 30 min each. Samples of 200 μL were collected from each treatment and stored at -80°C for insulin ELISA measurements. Dynamic tests of insulin secretion were carried out using a perifusion apparatus (Brandel Suprafusion SF-06, MD, USA) with a flow rate of 0.25 ml/min and sampling every 4 minutes. Samples from each fraction collected were analysed using insulin ELISA (Mercodia, Sweden). Following static and dynamic tests of insulin secretion, the SC-islets were collected, and the total insulin and DNA contents were analysed. Stimulated insulin secretion results are presented as fractional release of total insulin content after cell mass normalization using total DNA content.

### Cytotoxicity assays

A CD8^+^ T-cell line reactive to the HLA-A2-restricted peptide Influenza virus matrix protein MP_58-66_ (32% peptide-specific by tetramer staining) was thawed and rested for 3 h before use. Meanwhile, islet clusters were dissociated with TrypLE, stained with 1 μM CFSE or CellTrace Violet (CTV) and incubated for 2 h with 0.1 μM Influenza MP_58-66_ or peptide diluent, respectively. After washing, CFSE- and CTV-labeled SC-islets were mixed in equal numbers and cultured for 6 h in triplicate at 1x10^5^ cells/each per well in 96-well flat-bottom plates, alone or with increasing numbers of T cells (0.2x10^5^, 1x10^5^, 5x10^5^ and 10x10^5^), corresponding to effector-to-target (E:T) ratios of 1:5, 1:1, 5:1 and 10:1, respectively. After washing, cells were stained with Live/Dead Fixable Far Red (Thermo Fisher), antibodies against HLA-A, B, C (RRID: AB_2566151) and PD-L1 (RRID: AB_940368) and acquired on a BD LSRFortessa flow cytometer. SC-islets were analysed by FlowJo after gating on Live/Dead^−^ events and separation of CFSE^+^ (peptide-pulsed) and CTV^+^ (unpulsed) populations. Percent peptide-specific lysis at different E:T ratios is expressed as the ratio of live CFSE^+^/CTV^+^ cells normalized to the same ratio in wells containing SC-islet targets alone.

### Statistics

Data are collected from at least three independent differentiation experiments of 2 independent *TYK2* KO hiPSCs clones (C10 and C12). Blinding was applied for immunohistochemical quantification. Morphological data represents population-wide observation from independent differentiation experiments. Box and whiskers plots are presented as min to max showing all the points. Statistical methods used are described in each figure legend and individual method section. Briefly, Student’s unpaired two-tailed *t* test with Welch correction was used to compare differences between two groups while for more than two groups one-way ANOVA followed by Tukey’s test applied using Prism 8 software (GraphPad Software, La Jolla, CA). The results are presented as the mean ± S.D. unless otherwise mentioned. P-value < 0.05 were considered statistically significant (*P < 0.05; **P < 0.001; ***P < 0.0001).

### Study approval

The hiPSCs were generated and used according to the approval of the coordinating ethics committee of the Helsinki and Uusimaa Hospital District (no. 423/13/03/00/08).

Animal care and experiments were approved by National Animal Experiment Board in Finland (ESAVI/14852/2018).

## Author contributions

V.C. conceived and conceptualized the study, performed experiments, analysed data and wrote the first draft. H.I. carried out the experiments, standardized the differentiation, analysed data and participated in manuscript writing. C.H., F.V. carried out the T-cell cytotoxicity assays and analysed the data. R.P., O.P.D., L.G. performed Gene expression correlation analysis with fetal pancreas and adult islets RNAseq datasets and analysed the data. J.K. performed and analysed all the bulk and single cell RNAseq datasets. D.B. helped in generation of *TYK2* KO hiPSCs lines. J.S.V. and H.M. performed and analysed animal and differentiation experiments. T.B. and V.L. participated in the differentiation experiments and their analysis. I.A. provided and analysed human fetal RNAseq derived data. S.G. participated in microscopy and manuscript writing. R.M. conceptualized and supervised the T-cell cytotoxicity assays, analysed data, participated in manuscript writing and acquired funding. D.L.E. conceived and supervised the study, acquired funding and participated in manuscript writing. T.O. conceived and supervised the study, provided resources, acquired funding and wrote the manuscript. The assignment of authorship order for the first and second equal authors contribution is mainly chronological, detailed contributions described above.

## Acknowledgements

We gratefully acknowledge Dr. Fatoumata Samassa for help with cytotoxicity assays. We thank Jarkko Ustinov for the insulin and c-peptide ELISA measurements. S. Eurola, H. Grym and A. Laitinen are thanked for expert technical support. We thank FIMM Single Cell Analytics unit (supported by HiLIFE and Biocentre Finland) for single cell RNA sequencing services. We want to acknowledge the participants and investigators of the FinnGen study.

T.O. acknowledges the funding provided by the Academy of Finland (MetaStem Center of Excellence grant 312437), the Novo Nordisk Foundation and the Sigrid Juselius Foundation. R.M. acknowledges the support of the *Agence Nationale de la Recherche* (ANR-19-CE15- 0014-01) and the *Fondation pour la Recherche Medicale* (EQU20193007831). C.H. was funded by an *Année Recherche* fellowship of the Paris Saclay University. F.V. was funded by an international PhD fellowship of the IdEx Université de Paris. D.L.E. acknowledges the support of grants from the Welbio-FNRS (Fonds National de la Recherche Scientifique; WELBIO-CR-2019C-04), Belgium; the Innovate2CureType1 - Dutch Diabetes Research Foundation (DDRF), Holland; the Juvenile Diabetes Foundation (JDRF; 2-SRA-2019-834-S-B); the NIH (HIRN-CBDS) grant U01 DK127786, USA. D.L.E., T.O. and R.M. acknowledge support from the Innovative Medicines Initiative 2 Joint Undertaking under grant agreements No 115797 (INNODIA) and 945268 (INNODIA HARVEST), supported by the European Union’s Horizon 2020 research and innovation programme. These Joint Undertakings receive support from the Union’s Horizon 2020 research and innovation programme and “EFPIA”, “JDRF” and “The Leona M. and Harry B. Helmsley Charitable Trust”.

