## Supplemental Figures for "The type 1 diabetes gene *TYK2* regulates β-cell development and its responses to interferon-α"

Supplemental Figure 1

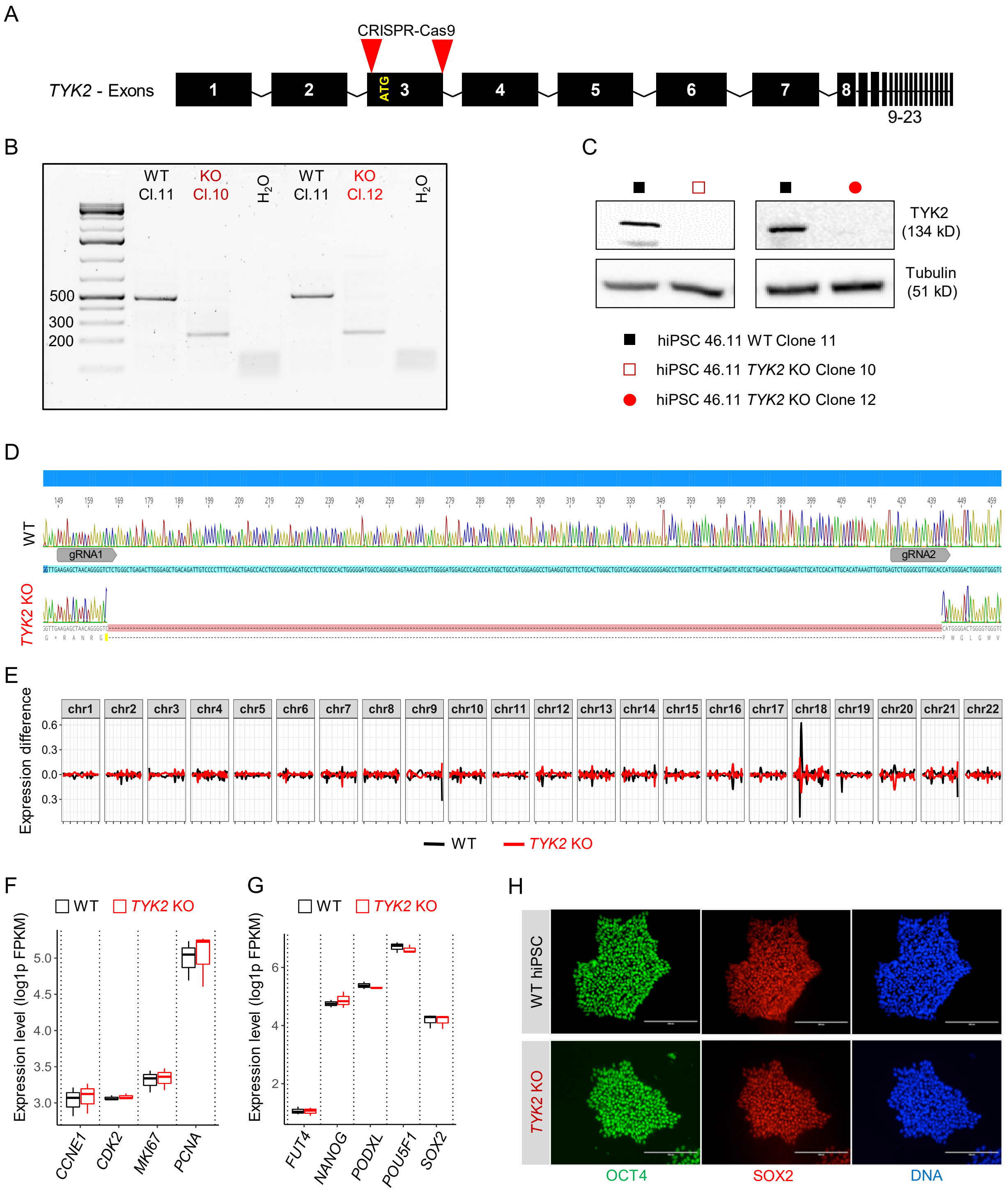

**Supplemental Figure1. Generation and characterization of *TYK2* knockout hiPSCs clones.**

**(A)** Schematic for the strategy to generate *TYK2* knockout by removing the ATG-containing-third exon in the *TYK2* gene of hiPSCs line HEL46.11. **(B)** A 504 bp PCR amplicon of *TYK2* gene showing a 277 bp deletion in KO clones. **(C)** Immunoblot based analysis for the expression of TYK2 protein in WT and *TYK2* KO hiPSC clones; tubulin protein expression shown for the loading control. **(D)** Sanger sequencing confirming the exact 277 bp deletion in the KO C10 clone, unedited WT clone C11 sequence also presented. **(E)** Global gene expression-based e-karyotyping analysis for WT and *TYK2* KO hiPSCs lines. **(F)** Boxplot showing the normalized FPKM values for the expression levels of proliferation markers, *CCNE1*, *CDK2*, *MKI67*, *PCNA* and **(G)** pluripotency factors, *FUT4*, *NANOG*, *PODXL*, *POU5FI*, *SOX2* utilizing whole transcriptome data of WT and *TYK2* KO hiPSCs line. **(H)** Representative image of immunocytochemistry analysis for the expression of pluripotency factor OCT4 and SOX2 in WT and *TYK2* KO hiPSCs. Scale bar = 200 μm.

### Supplemental Figure 2

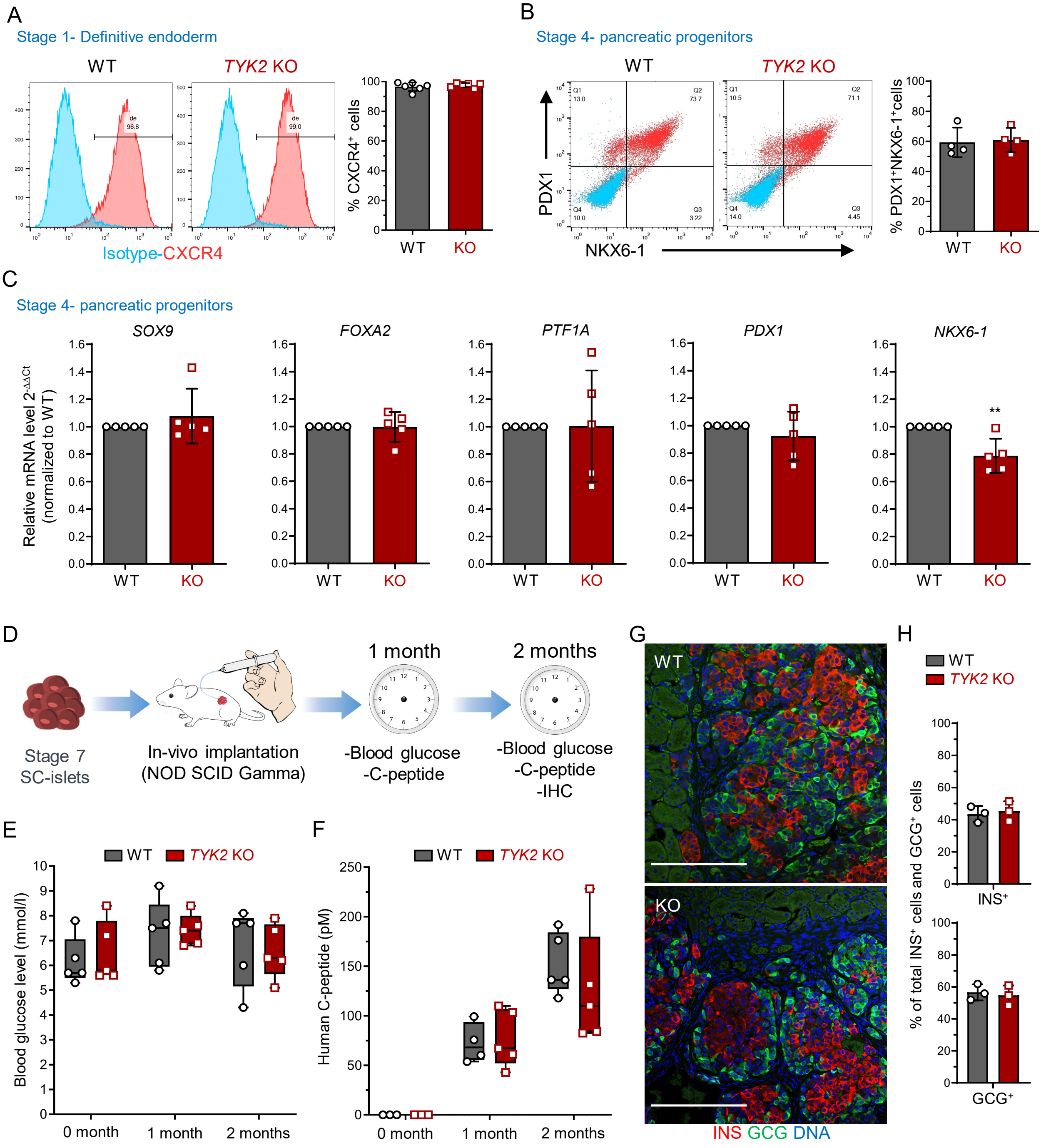

**Supplemental Figure 2.** ***In vitro*** **and** ***In vivo*** **characterization of WT and** ***TYK2*** **KO hiPSCs pancreatic differentiation.**

Flow cytometry analysis for the expression of **(A)** CXCR4$^+$ definitive endoderm cells at the end of stage-1 and **(B)** PDX1$^+$ and NKX6-1$^+$ double positive cells at the end of stage-4 differentiation in WT and *TYK2* KO cells. **(C)** Expression of stage-4 pancreatic progenitor markers *SOX9*, *FOXA2, PTF1A, PDX1* and *NKX6-1* with qRT-PCR. Two-tailed unpaired t-test was performed to determine the significance levels (n = 4 - 6). **(D)** Schematic of *in vivo* implantation study in NOD/SCID-γ mice. S7 WT and *TYK2* KO SC-islets were implanted in the mice (n = 4-5, for each genotype). **(E)** Mouse blood glucose levels measured at day 0-, 1- and 2-months following implantation; **(F)** Human C-peptide levels measured in the mice sera at 0-, 1- and 2-months following implantation. Box and whiskers plots showing min to max with all the points. **(G-H)** Immunohistochemistry of grafts after 2 months of implantation showing insulin (INS) and glucagon (GCG) and their quantification. Two-tailed unpaired t-test were performed to determine the significance levels. Data are mean ± S.D. **p < 0.01. Scale bar = 200 μm.

Supplemental Figure 3

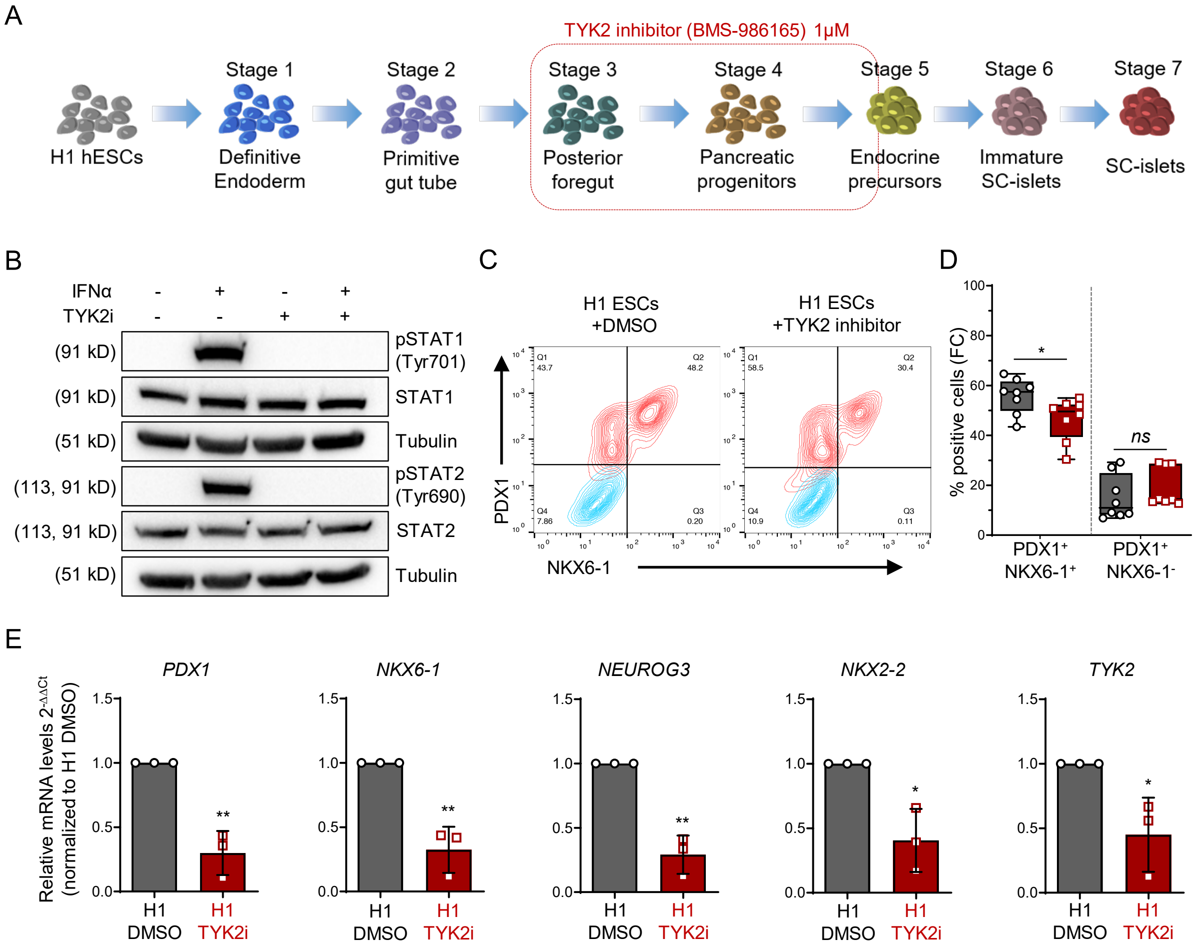

**Supplemental Figure 3. TYK2 inhibitor (TYK2i; BMS-986165) treatment during early pancreatic differentiation compromised endocrine precursors formation in hESCs line H1.** **(A)** Human ES cell-line H1 differentiated with TYK2i as per schematic of the experiment plan. **(B)** Phosphorylation of STAT1 and STAT2 following 30 min IFNα treatments in DMSO or TYK2i treated differentiated cells (n = 3) at stage 5. **(C-D)** Representative contour plot of flow cytometry analysis following TYK2i treatment for the determination of $PDX1^{+}NKX6\text{-}1^{+}$ and $PDX1^{+}NKX6\text{-}1^{-}$ cells and their quantification (n = 8). Box and whiskers plot showing min to max with all the points. **(E)** Expression of *PDX1*, *NKX6-1*, *NEUROG3*, *NKX2-2* and *TYK2* with qRT-PCR (n = 3). Two-tailed unpaired t-test was performed to determine the significance levels. Data are mean ± S.D. *$p < 0.05$; **$p < 0.01$.

Supplemental Figure 4

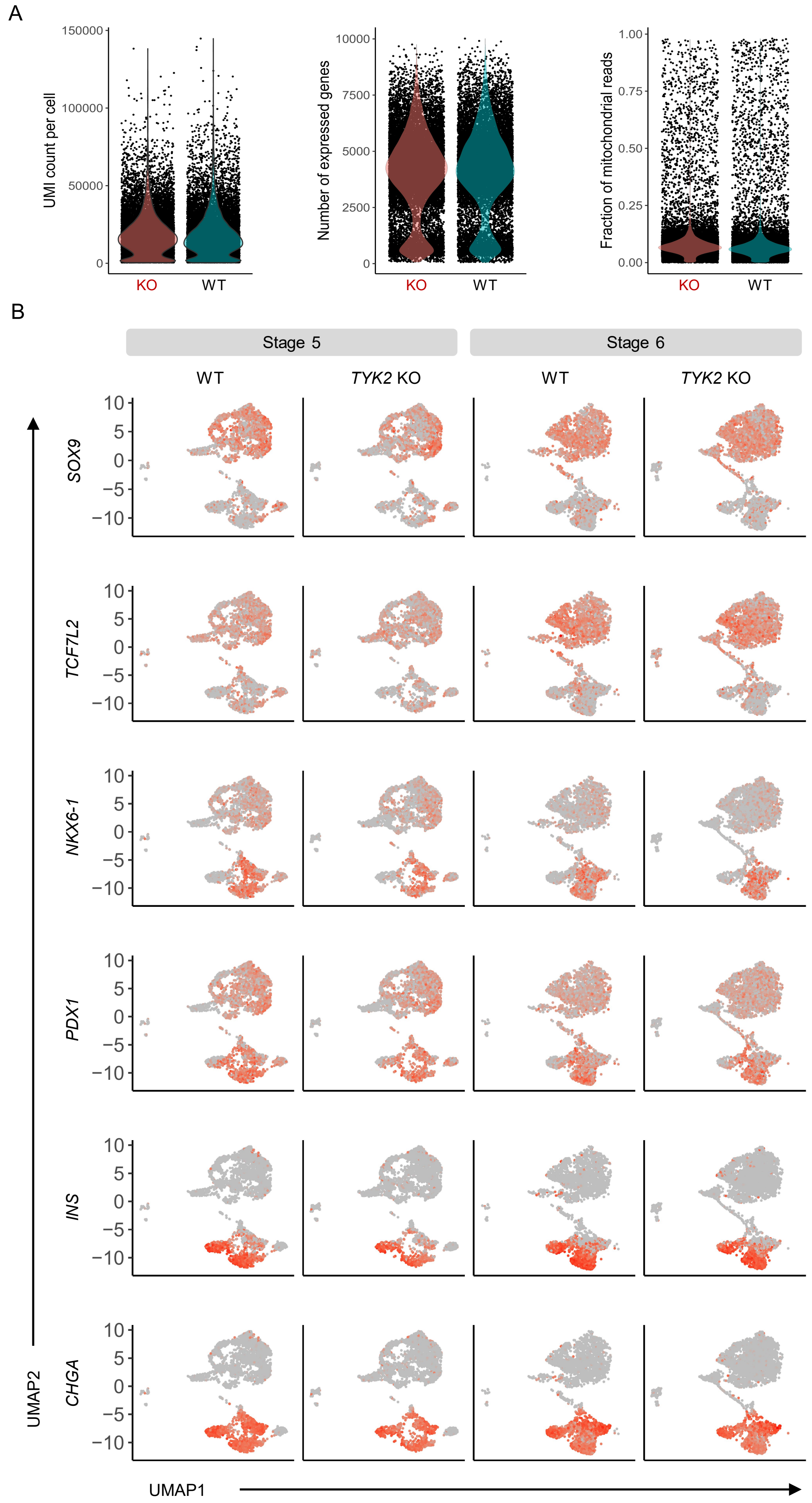

**Supplemental Figure 4. Quality assessment of single cell RNA sequencing analysis.**

**(A)** Average unique molecular identifier (UMI) counts, number of expressed genes and mitochondrial reads per cell were plotted for the scRNA-seq analysis. Cells with less than 5000 UMI counts or 1700 expressed genes or with unusually high levels of mitochondrial reads (>20% of counts) were excluded. **(B)** Feature plot showing the expression pattern of *SOX9*, *TCF7L2*, *NKX6-1*, *PDX1*, *INS* and *CHGA* for stages 5 and 6 in WT and *TYK2* KO scRNA-seq samples.

Supplemental Figure 5

A

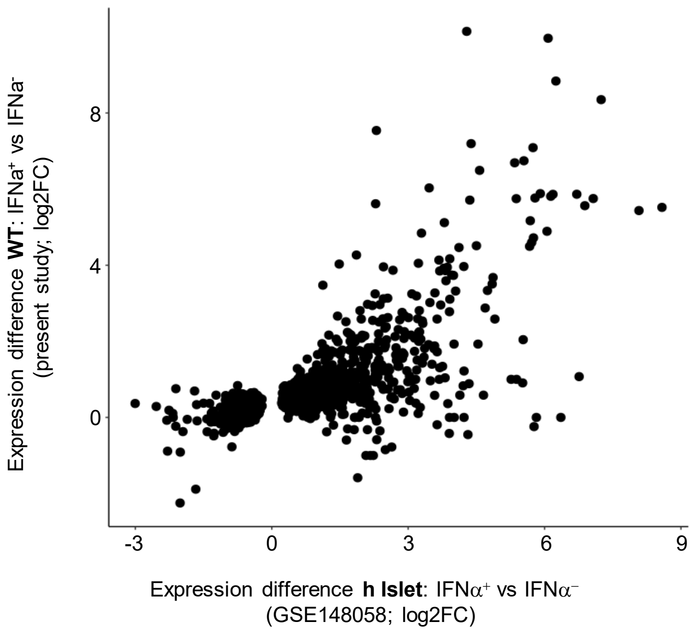


B

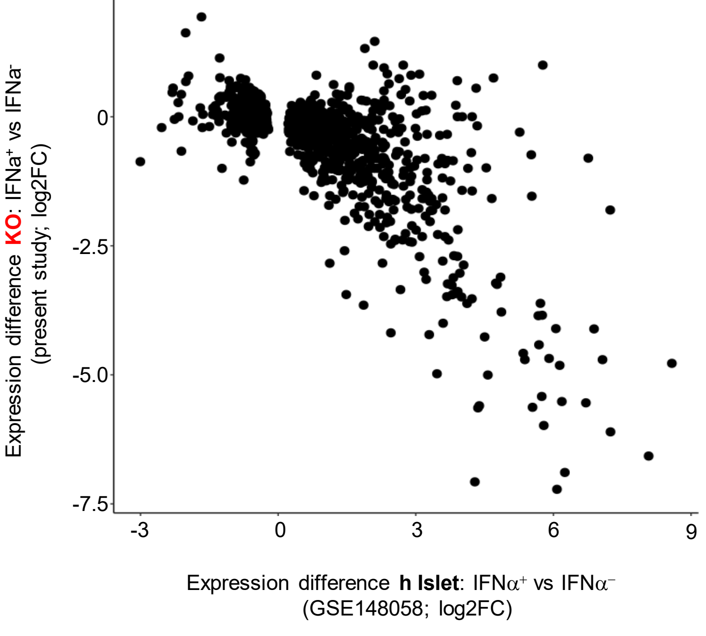

**Supplemental Figure 5. SC-islets exposed to IFNα show similar global transcriptomic changes compared to human islet samples under similar treatment.**

Log2FC of the differentially expressed genes from a previous study (GSE148058) of human islets treated with or without IFNα (-/+) for 18h at X-axis versus the values of **(A)** WT or **(B)** *TYK2* KO SC-islets, treated for 24h with IFNα in the present study at Y-axis, respectively.

Supplemental Figure 6

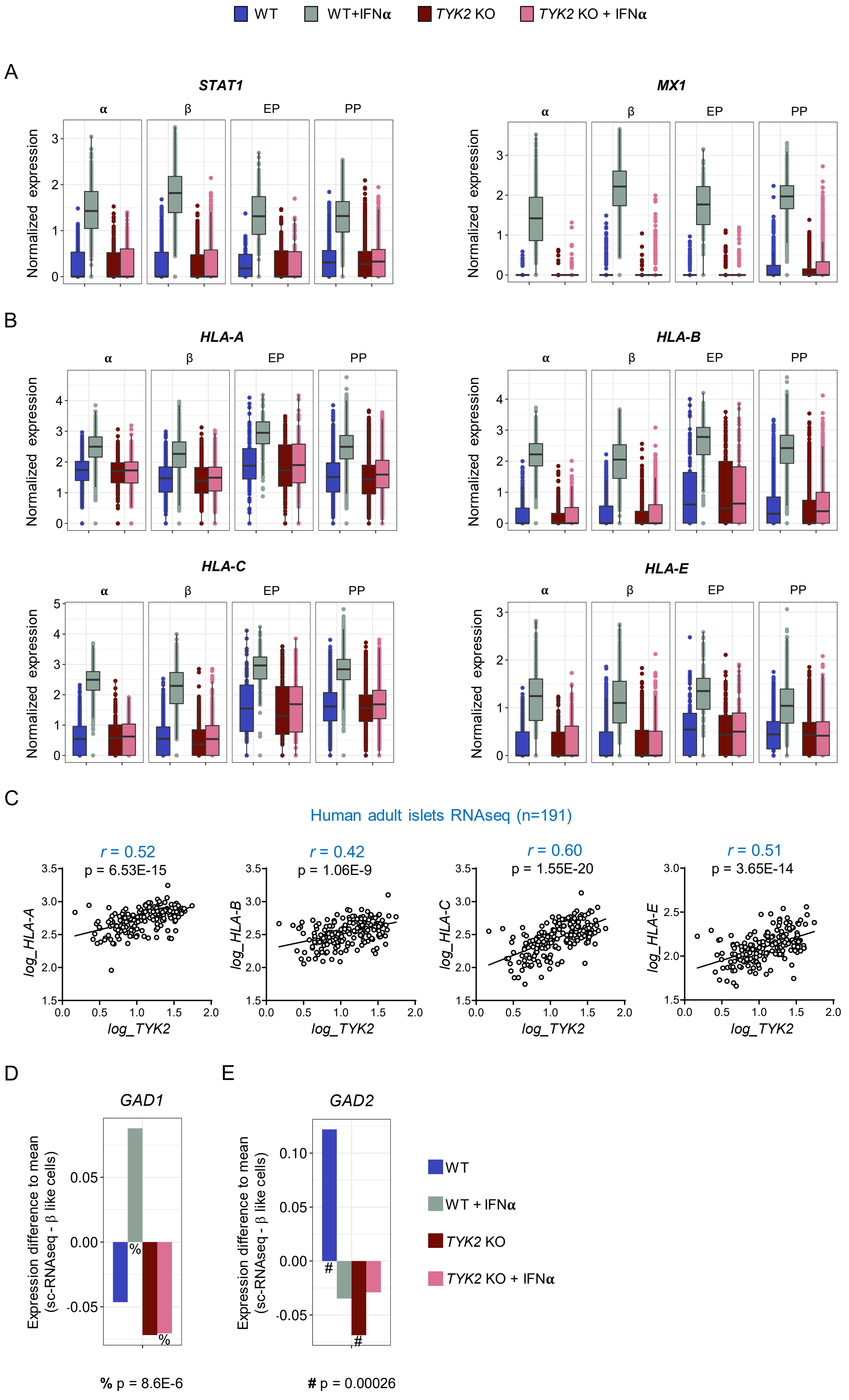

**Supplemental Figure 6. TYK2 regulates the expression of IFNα stimulated genes and autoantigen (GAD1 and GAD2) in SC-islets.**

Single cell transcriptomics performed on the WT and *TYK2* KO S6 SC-islets following 24h -/+ IFNα treatment. Condensed Boxplot showing the normalized expression of IFNα stimulated genes **(A)** *SATA1* and *MX1*, **(B)** *HLA-A*, *HLA-B*, *HLA-C* and *HLA-E* in the clusters of α-, β-, endocrine precursor (EP)- and pancreatic progenitor (PP)- like cells. **(C)** Gene expression correlation between *TYK2* and *HLA-A, HLA-B, HLA-C, HLA-E* with human islets RNA-seq samples (n = 191). Pearson correlations *r* and significance levels *p* are also indicated in the panel. **(D-E)** The Bar plots showing the expression pattern for the autoantigens GAD1 and GAD2 in different genotypes indicated by color codes.

Supplemental Figure 7

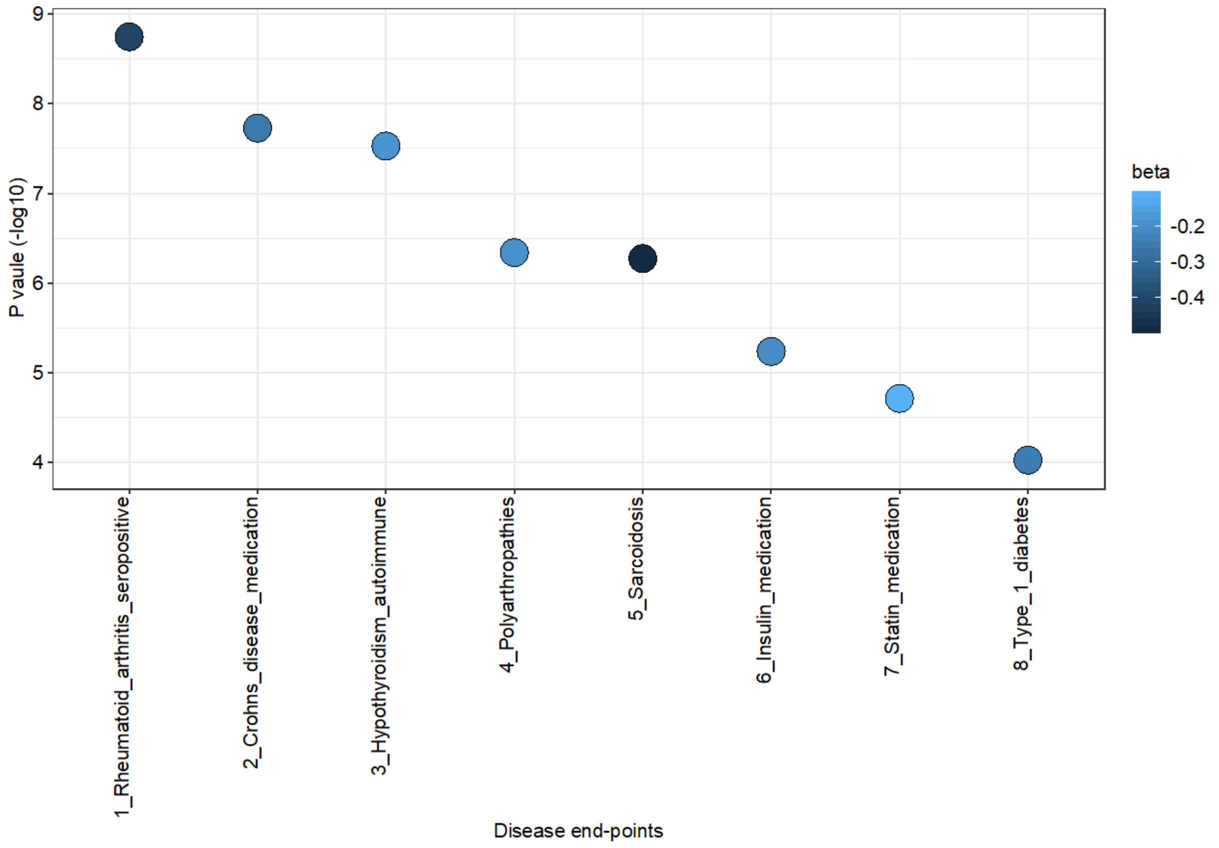

**Supplemental Figure 7. TYK2$^{P1104A}$ (rs34536443) protects against autoimmune or auto-inflammatory diseases in Finnish population.**

The association analysis of TYK2$^{P1104A}$ with 2803 clinical endpoints obtained from electronic health record data (ICD codes and drug purchase data) of 218,792 Finnish individuals. The figure depicts the effect size (beta) and the corresponding significance of association for the top associated unique clinical endpoints (Rheumatoid arthritis- seropositive, cases = 4,594, controls = 214,196; Crohn's disease medication, cases = 9,669, controls = 209123; Hypothyroidism (autoimmune), cases = 22,997, controls = 175,475; Polyarthropathies, cases = 15,246, controls = 147,221; Sarcoidosis, cases = 2,046 , controls = 215,712; Insulin medication, cases =11,590, controls = 185,895 ; Statin medication, cases = 68,782, controls = 150,010 ; Type-1-diabetes, cases = 5,928, controls = 183,185). The association analysis was done using mixed model logistic regression adjusted for age, sex, 10 PCs and genotype batch.
